# Progressive lengthening of 3′ untranslated regions of mRNAs by alternative cleavage and polyadenylation in cellular senescence of mouse embryonic fibroblasts

**DOI:** 10.1101/031302

**Authors:** Miao Han, Guoliang Lv, Hongbo Nie, Ting Shen, Yichi Niu, Xueping Li, Meng Chen, Xia Zheng, Wei Li, Chen Ding, Gang Wei, Jun Gu, Xiao-Li Tian, Yufang Zheng, Xinhua Liu, Jinfeng Hu, Wei Tao, Ting Ni

## Abstract

**Background:** Cellular senescence has historically been viewed as an irreversible cell cycle arrest that acts to prevent cancer. Recent discoveries demonstrated that cellular senescence also played a vital role in normal embryonic development, tissue renewal and senescence-related diseases. Alternative cleavage and polyadenylation (APA) is an important layer of post-transcriptional regulation, which has been found playing an essential role in development, activation of immune cells and cancer progression. However, the role of APA in the process of cellular senescence remains unclear.

**Materials and Methods:** We applied high-throughput paired-end polyadenylation sequencing (PA-seq) and strand-specific RNA-seq sequencing technologies, combined systematic bioinformatics analyses and experimental validation to investigate APA regulation in different passages of mouse embryonic fibroblasts (MEFs) and in aortic vascular smooth muscle cells of rats (VSMCs) with different ages.

**Results:** Based on PA-seq, we found that genes in senescent cells tended to use distal pA sites and an independent bioinformatics analysis for RNA-seq drew the same conclusion. In consistent with these global results, both the number of genes significantly preferred to use distal pAs in senescent MEFs and VSMCs were significantly higher than genes tended to use proximal pAs. Interestingly, the expression levels of genes preferred to use distal pAs in senescent MFEs and VSMCs tended to decrease, while genes with single pAs did not show such trend. More importantly, genes preferred to use distal pAs in senescent MFEs and VSMCs were both enriched in common senescence-related pathways, including ubiqutin mediated proteolysis, regulation of actin cytoskeleton, cell cycle and wnt signaling pathway. By cis-elements analyses, we found that the longer 3′ UTRs of the genes tended to use distal pAs progressively can introduce more conserved binding sites of senescence-related miRNAs and RBPs. Furthermore, 375 genes with progressive 3′ UTR lengthening during MEF senescence tended to use more strong and conserved polyadenylation signal (PAS) around distal pA sites and this was accompanied the observation that expression level of core factors involved in cleavage and polyadenylation complex was decreased.

**Conclusions:** Our finding that genes preferred distal pAs in senescent mouse and rat cells provide new insights for aging cells’ posttranscriptional gene regulation in the view of alternative polyadenylation given senescence response was thought to be a tumor suppression mechanism and more genes tended to use proximal pAs in cancer cells. In short, APA was a hidden layer of post-transcriptional gene expression regulation involved in cellular senescence.

## INTRODUCTION

Cellular senescence was originally described as a process that limited the proliferation of normal human fibroblasts in culture, and caused by the loss of telomeres after extensive proliferation in the absence of endogenous telomerase activity [1, 2]. In fact, besides telomere erosion, many stimuli and stress could also cause cellular senescence, including DNA double strand breaks, strong mitogenic signals, oxidative stress, loss of the PTEN tumor suppressor and ectopic expression of the cyclin-dependent kinase inhibitors (CDKIs) [3]. Morphological changes and molecular markers of senescent cells were identified in the past decades [4, 5]. These features include a flattened and enlarged cell morphology, absence of proliferative markers, senescence-associated β-galactosidase activity (SA-β-gal), expression of tumor suppressors, cell cycle inhibitors, and also DNA damage markers, including nuclear foci of constitutive heterochromatin and prominent secretion of growth factor, proteases, cytokines and other factors that have potent autocrine and paracrine activities [3, 6]. Noteworthy, no hallmark of senescence identified thus far is entirely specific to the senescent state, and not all senescent cells express all possible senescence markers [3].

Cellular senescence was generally viewed as an important mechanism for preventing the growth of cells at risk for neoplastic transformation [7, 8]. In recent years, it has become apparent that cellular senescence can also be involved in multiple physiological and pathological processes, such as normal embryonic development and tissue damage, suggesting that cellular senescence plays both beneficial and detrimental roles [6]. In younger individuals, cellular senescence blocks proliferation of impaired cells, thus preventing them from transforming into cancer cells and leads to anti-cancer and anti-aging benefits [6, 9]. However, when people get old, a growing number of cells enter senescence, which causes decreased tissue function, elevated inflammation and exhaustion of stem cells, thus become pro-aging [6, 9]

Cellular senescence can be generally divided into replicative senescence and stress-induced premature senescence (SIPS) [6]. Primary cells including human diploid fibroblasts (HDFs) and mouse embryonic fibroblasts (MEFs) that cultured *in vitro* are often used as model systems to investigate the mechanism of replicative senescence[10, 11]. It is noteworthy that MEFs enter replicative senescence at regular culture condition where oxygen level is high, the senescence won’t be established if given low oxygen tension[12–14]. Similarly, HDFs under more physiological O_2_ concentration (3%) are achieved more numbers of population doublings than the O_2_ concentration of air (20%)[15]. Further studies demonstrated that DNA damage response (DDR) contributes to *in vitro* replicative senescence for both HDFs and MEFs [12, 15]. Interestingly, MEFs have significant less numbers of population doublings (PD) than HDFs *in vitro,* making MEFs a time-saving model for investigation of replicative senescence.

Several signaling pathways have been functionally involved in the progression of cellular senescence and *in vivo* ageing including telomere attrition, epigenetic alternations, deregulated nutrient-sensing, mitochondrial dysfunction, altered intercellular communication, exhaustion of stem cells [9]. Notably, most of these pathways have not been evolved as direct regulators of ageing, such as nutrient signaling is critical in promoting growth effects during embryogenesis and early development[16]. Intensive studies have attempted, through high-throughput genome-wide transcriptomics, to identify gene expression signatures that define cellular senescence. Many studies have found that dramatic changes in transcriptomes occur accompany the dramatic phenotype changes of senescent cells [10, 17–19]. Kim et al. profiled cellular senescence phenotype and mRNA expression patterns during replicative senescence in human diploid fibroblasts [10]. Interestingly, they found that the gene expression modules governing each stage supported the development of the associated senescence phenotypes [10]. Furthermore, Mazin et al. found widespread splicing changes during human brain development and aging [19]. Other studies also suggested that modification of mRNA processing may be a feature of aging [18, 20, 21]. However, our knowledge about regulation of the transcriptome in cellular senescence is still limited. RNA processing is a multi-step but tightly regulated pathway which includes transcription, splicing, editing, transport, stability, localization, translation and degradation [22]. What is more, co- and post-transcriptional regulation of gene expression is very complex and multifaceted, covering the complete RNA lifecycle from biogenesis to decay [23]. Increasing evidence supports that post-transcriptional regulation plays important roles during normal brain aging and cellular senescence through interacting between microRNAs and 3′ untranslated region (3′ UTR) of target genes, or interacting between RBPs (RNA binding proteins) and 3′ UTR of cyclin-dependent kinase inhibitors, such as p16 [24, 25]. These results raised the possibility that changes of 3′ UTR would also contribute to cellular senescence and/or aging.

Cleavage and polyadenylation of nascent RNA is essential for maturation of the vast majority of eukaryotic mRNAs and determines the length of in the 3′ UTR. Such process requires several cis-acting RNA elements and several dozens of core and auxiliary polypeptides [26]. The key cis-element that defines cleavage is a 6 nucleotide (nt) motif called the poly(A) signal (PAS), the canonical form of which is AAUAAA. However, it can adopt more than ten variants thereof and is mostly located in 40nt upstream of the cleavage site [26]. In addition to the PAS, additional U-rich sequence elements located upstream as well as U-rich and GU-rich sequence elements located downstream of the cleavage site also can enhance the efficiency of the 3′ end processing reaction [26]. The 3′ end-processing machinery, contained several sub-complexes, including cleavage and polyadenylation specificity factor (CPSF), cleavage stimulation factor (CstF), cleavage factor I and II (CFI and CFII), as well as additional accessory such as Poly(A) polymerase (PAP), nuclear poly(A) binding protein 1 (PABPN1), Symplekin and RNA polymerase II (RNAAP II), is responsible for the cleavage and polyadenylation reaction[27]. With the surge of high-throughput sequencing technologies, genomic studies in the past few years have indicated that most eukaryotic mRNA genes have multiple polyadenylation sites (pAs) and thus multiple 3′ UTRs, caused by alternative cleavage and polyadenylation (APA) [27]. For examples, about 72% genes in *Saccharomyces cerevisiae,* 70% in Arabidopsis, 30% in *C. elegans,* 43% in zebrafish, 79% in mouse, and 69% in human undergo APA [28–34]. In addition, 66% of mouse long noncoding RNAs have also been found to have APA, and a large amount of APA events in long noncoding RNAs have also been found in different human tissues [29, 35].

Alternative pAs can reside in the 3′-most exon or upstream regions, leading to multiple mRNA isoforms that contain different coding sequences, 3′ untranslated regions (3′ UTRs) or both [26]. In fact, more than a hundred different conserved sequences have been identified in mRNA 3′ UTRs alone and can be considered as putative regulatory elements [36]. These sequences are divided into two main classes depending on the nature of their trans-acting factors which can be either microRNAs (miRNA) or RNA binding proteins (RBPs). Both miRNA and RBPs have been reported to control gene expression by modulating the translational efficiency or degradation of their mRNA targets while the specific control of mRNA subcellular localization relies more exclusively on RBP interactions [37]. Therefore, APA can play a significant role in mRNA metabolism, such as stability, translation, and subcellular localization, through changing the interactions between miRNAs and/or RBPs with the 3′ UTRs region by controlling the length of the 3′ UTRs [38, 39].

Furthermore, dynamic regulation of 3′ UTRs by APA has been reported in different tissue types, development and cellular differentiation, cell proliferation, cell reprogramming and cancer cell transformation [31, 35, 40–47]. Especially, a large scale analysis of five mammals showed that pA usage is more conserved in the same tissue across species than in different tissues within the same species, highlighting an essential role of APA in establishing tissue-specific gene expression profiles [28]. Sanberg et al. first reported the link between APA and cell proliferation, indicating fast growing cells favored proximal pA sites [43]. Consistent with the association between cell proliferation and APA, cancer cells also preferred expression of mRNA isoforms with shorter connection 3′ UTRs compared with normal cells[44, 48, 49]. Progressive 3′ UTR lengthening coupled with weakening of mRNA polyadenylation activity was observed during mouse embryonic development [42]. Conversely, 3′ UTRs become shorter and the expression of mRNAs encoding cleavage and polyadenylation (C/P) factors increased during reprogramming of somatic cells from several different cell types [47]. In addition, the regulation of APA is dictated by a combination of several features, including variations in the abundance or activity of *trans-acting* factors such as core 3′ processing factors, tissue-specific RBPs, as well as splicing and transcription factors, combinations of *cis*-acting RNA elements, nucleosome positioning, DNA methylation and histone post-translational modifications [27, 50]. To sum up, alternative cleavage and polyadenylation of transcripts from the same gene enables cell-type- or condition-specific expression of 3′ UTR isoforms, there by adding another layer of post-transcriptional regulation. However, the extent to which differential expression of these isoforms is used to regulate mRNA and protein levels in cellular senescence is completely unclear.

In this study, we applied our published polyadenylation sequencing (PA-seq) and strand-specific RNA-seq sequencing technologies, followed by systematic bioinformatics analyses and experiments to investigate APA regulation in different passages of mouse embryonic fibroblasts (MEFs) and in aortic vascular smooth muscle cells of rats (VSMCs) with different ages. Based on PA-seq and RNA-seq data, this study found that in senescent cells, many genes tended to use distal pA sites, together with decreased expression levels and also preferred to enriched in common senescence-related pathways. Interestingly, we found that genes preferred to use distal pAs in cellular senescence tended to use proximal pAs in cancer cells [48]. This was consistent with the previous knowledge that senescence response was a potent tumor suppressive mechanism [8]. We further found that global 3′ UTR lengthening in senescent cells might be explained by decreased expression of cleavage and polyadenylation related factors. These above results provide first evidence to suggest that APA is a hidden layer of gene’s post-transcriptional regulation during cellular senescence.

## RESULTS

### Establishment of replicative senescence model for MEFs

To uncover the role of alternative polyadenylation (APA) in cellular senescence, we isolated and continuously subcultured mouse embryonic fibroblasts (MEFs) to serve as a typical replicative senescence model [11–14]. Population doubling curve showed that MEF cells underwent decreased growth rate (Figure 1A), a traditional marker for replicative senescence. Cells with population doubling time (PD) 6, 8, 10 and 11 were further analyzed for additional senescence marks. Mki67, an indicator for cell vitality and also a robust assessment of cellular senescence [51], showed decreased expression level that determined by quantitative reverse transcription PCR (qRT-PCR) (Figure 1B). Flow cytometry revealed reduced percentage of S phase cells from PD6 to PD11 (Figure 1C). P16, a well-known cyclin-dependent kinase inhibitor that involved in multiple cellular senescence types, exhibited elevated expression at both RNA (Figure 1D) and protein level (Figure 1E). Senescence-associated β-galactosidase activity (SA-β-gal) also increased from PD6 to PD11 (Figure 1F). All these above evidences support that MEF cells from PD6 to PD11 underwent replicative senescence and can be further applied to investigate the potential regulation of APA.

**Figure1.**
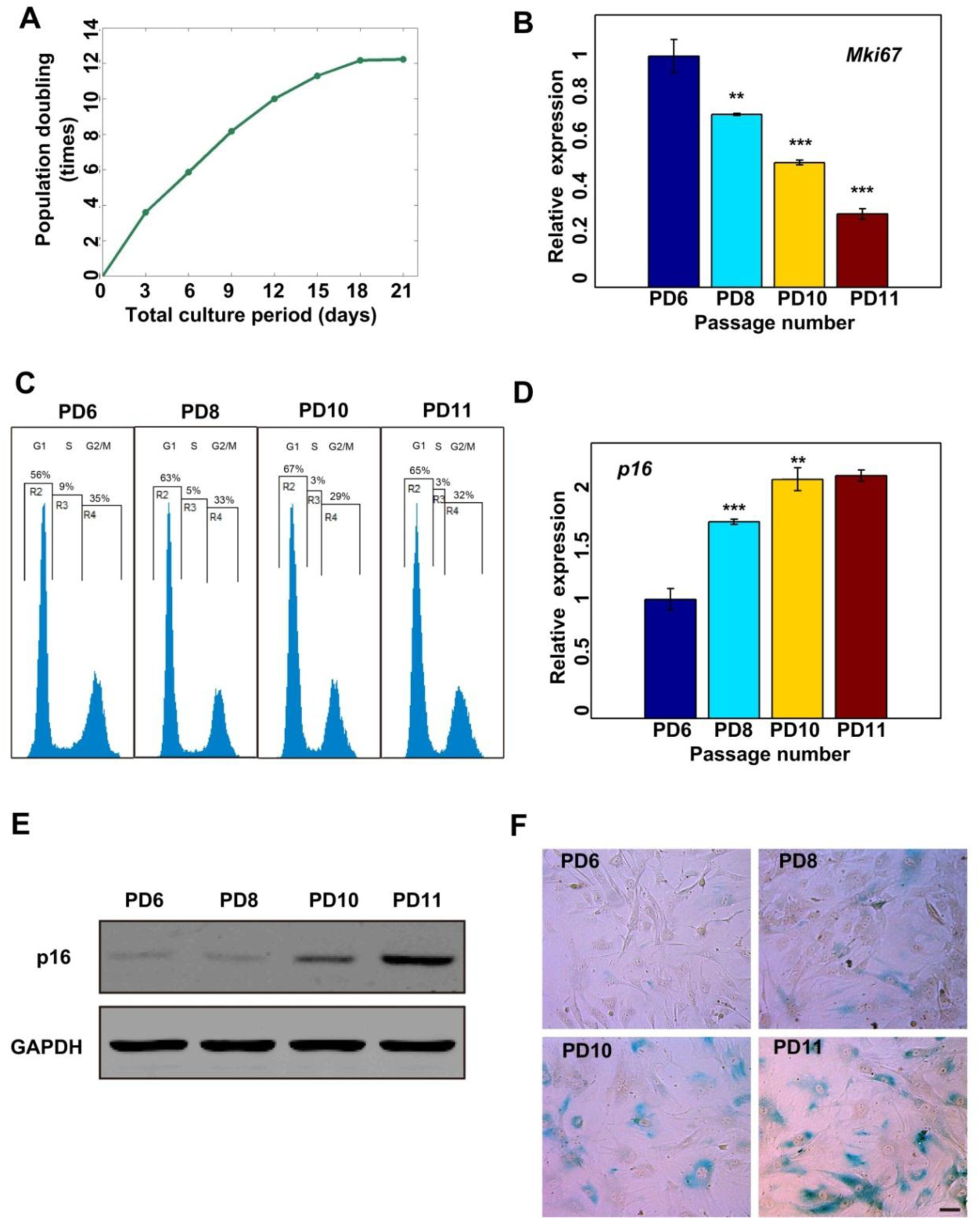
Establishment of a replicative senescence model for MEFs. (A) MEFs were continuously subcultured in DMEM medium containing 10% FBS (fetal bovine serum) and their population doubling numbers (passaged numbers of the cells obtained by sequential population doubling) were recorded. (B) Detection expression levels of *Mki67* in PD6, PD8, PD10 and PD11 passages of MEFs by qRT-PCR. (C) MEFs form PD6, PD8, PD10, PD11 passages were subjected to FACS (Fluorescence-activated cell sorting) analysis. (D) Expression levels of p16 (encoded by *Cdkn2a*) in PD6, PD8, PD10 and PD11 passages of MEFs were detected by RT-qPCR. (E) Detection of p16 protein levels in PD6, PD8, PD10 and PD11 passages of MEFs by Western Blot. (F) MEFs from PD6, PD8, PD10 and PD11 passages were subjected to SA-β-gal (senescence-associated β-galactosidase) staining. (***) *P* <0.001, (**) *P* <0.01 and (*) *P* <0.05, two tailed Student’s t-test.

### Prevalent APA in protein-coding and noncoding transcripts in MEFs

To explore the potential role of APA in replicative senescence, we performed our published PA-seq protocol [35] to map the 3′ end of mRNAs in 4 different passages of MEF cells (PD6, PD8, PD10, PD11), which showed multiple senescence markers in later population doubling, including slower growth rate and elevated SA-β-gal activity (Figure 1). Previous studies have demonstrated that PA-seq can not only accurately and comprehensively identify poly(A) sites (pAs) in mRNAs and noncoding RNAs, but also provide quantitative information on the relative abundance of polyadenylated RNAs [35]. As an independent survey of global APA regulation in cellular senescence, strand-specific RNA-seq for the same four MEF samples (PD6, PD8, PD10, PD11) was also carried out, followed by a dedicated data analysis pipeline using RUD index developed by Tian’s lab [52]. In order to evaluate the influence of cell cycle phase, which also associated with cellular senescence, to APA regulation, we investigated the APA of cells in G0 phase through serum starvation of PD6 cell (the sample is called G0 hereafter). These above experimental designs enable us to address how widespread is the APA events and what’s the global changes of APA during replicative senescence.

In total, we obtained ~54 million strand-specific paired-end 101-mer reads for PA-seq. Out of these, ~63% (about ~34 millions) can be uniquely mapped to the mouse reference genome (see details in Supplementary Table S1). We applied F-seq, a feature density estimator [53] to compute PA cluster based on uniquely mapped reads. The PA-seq data from 4 samples (PD6, PD8, PD10, PD11) and also the G0 stage were combined so that a unified peak-calling scheme can be applied. A total of 29,144 PA clusters were obtained after applying the following two criteria: (i) To filter out any potential internal priming events during reverse transcription, we removed PA clusters with 15 ‘A’ in the 20 nucleotides region downstream of peak mode (the most frequent position in the cluster) or with continuous 6 ‘A’ in the downstream of peak mode; (ii) Any peak with less than 20 tags was then removed to obtain reliable PA clusters. We further evaluated the genomic distribution of these PA clusters. Among these 29,144 high quality peaks, majority (26,905 or 92.3%) were located within the genomic region of RefSeq gene annotation (also include the 250 bp upstream and the 5000 bp downstream of the gene body, which may be the potential region of the promoter or the distal pAs of the genes), suggesting most of the PA peaks belongs to annotated genes. The remaining 2,239 (or 7.7%) PA clusters were mapped to the intergenic regions, suggesting a considerable number of novel intergenic transcripts also contain poly(A) tail. In order to focus the analysis on robustly expressed APA events, we kept clusters that accounted for ≥ 10% of all the tags within respective genes in any sample. If a lowly expressed PA cluster contains > 5% of all the tags within respective genes in the majority of samples (≥ 80%), the peak was also kept. By such filtering of lowly expressed peaks, we got 18,175 and 464 high confident PA clusters in coding genes and noncoding RNAs, respectively (Figure2A, Supplementary Table S2). 65.8% of these 18,639 high confident PA clusters are included in the PolyA_DB 2 (Supplemental Figure S1A), which contains pA sites (pAs) for genes in several vertebrate species, including human, mouse, rat, chicken and zebrafish, using alignment between cDNA/ESTs and genome sequences [54]. We next examined the genomic distribution of identified PA clusters or sites located in protein-coding genes. 42% of the PA clusters are overlapped with known polyadenylation sites (Known pA), while 41% and 10% clusters are located in the annotated and extended 3′ UTR regions, respectively (Figure 2B). In addition, a considerable proportion (4%) of the PA clusters fall into introns, and a small subset of the PA clusters is mapped to exons (2%) as well as the upstream regions of the transcripts (<1%). Together, our result support the notion that 3′-end formation in the MEF transcriptomes is much more complex than previously appreciated as in the human transcriptomes [33, 35, 55]. We further evaluated the tag abundance in the top three categories of PA clusters (Known pA, 3′ UTR and extended 3′ UTR in Figure 2C). Consistent with previous knowledge that annotated pAs have the highest expression, 56% of all the tags covering the known polyadenylation sites (Known pA in Figure 2C). Further analysis showed that the overall tag number distribution in known pA site also tended to be higher compared to those in 3′ UTR or extended 3′ UTR regions (Supplemental Figure S1B). In addition, the median number of covered tags for pAs located in tandem 3′ UTR is significantly higher than in both exon and intron regions, respectively (*P*<0.001 and *P*<0.001, respectively, Mann-Whitney U test, see Supplemental Figure S1C). These above results indicate that pAs located in known pA sites, 3′ UTR and extended 3′ UTR regions are three major groups in MEF cells.

**Figure2.**
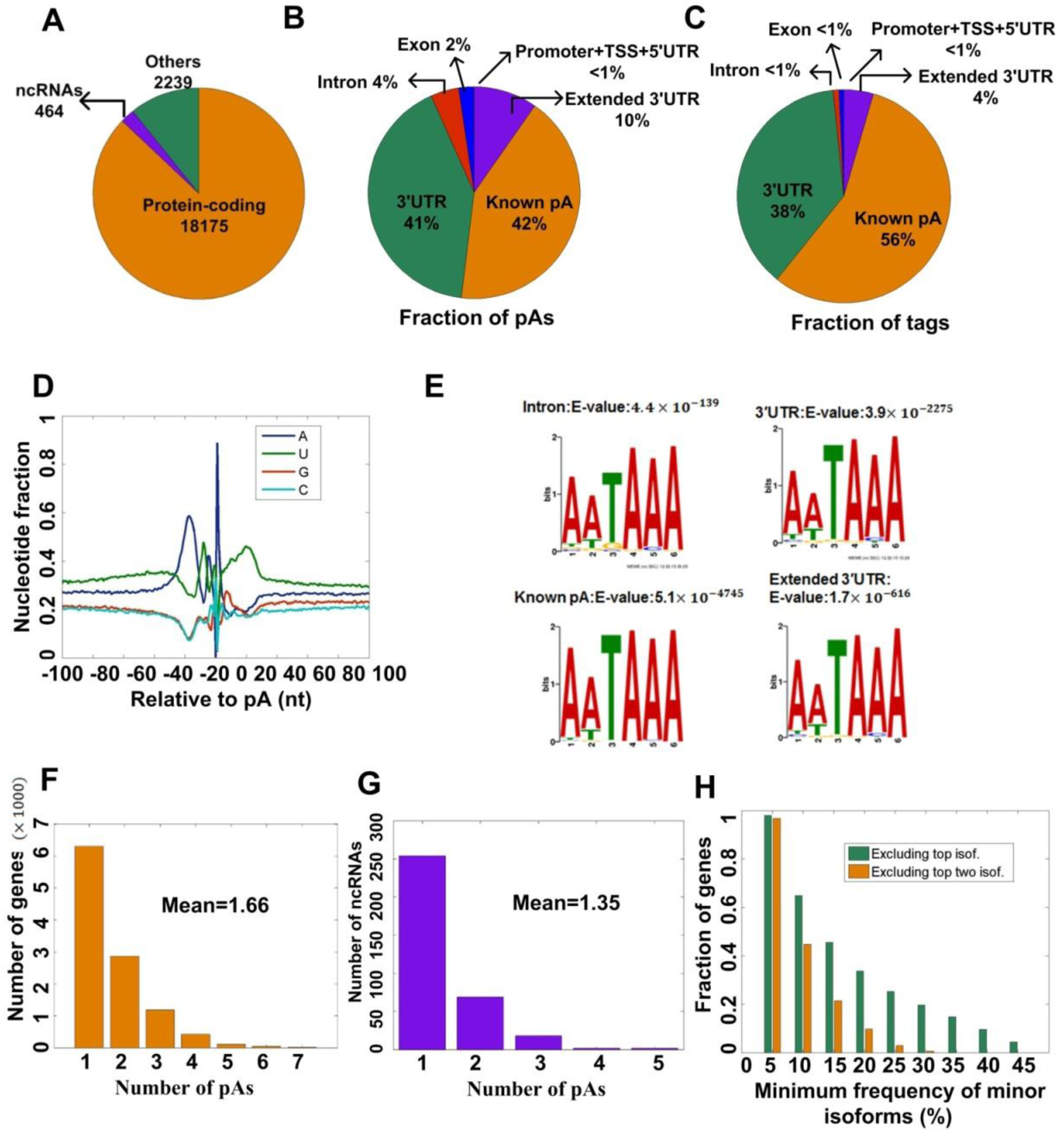
Summary of pAs in MEFs identified by PA-seq. (A) Gene categories of pAs (poly(A) sites) based on RefSeq annotation. (B) The relative locations of the identified pAs in respective genes. The pAs classification scheme is described in materials and methods. (C) The relative locations of the identified pA tags in respective genes. (D) Nucleotide sequence composition around the pAs identified in this study. (E) MEME program identifies the canonical PAS (poly(A) signal) motif AATAAA with a very significant E-value from the upstream (-40bp) of the pAs located in intron, 3′ UTRs, known pAs and extended 3′ UTR regions. (F) The number of genes with different number of pAs. (G) The number of ncRNAs with different number of pAs. (H) Fraction of genes with the minimum frequency of minor 3′ UTR isoforms exceeding given thresholds (0%, 5%, 15%, 20%, 25%, 30%, 35% 40% and 45%). Frequencies of minor isoforms are calculated by excluding the top isoform (green bars) and the top two isoforms (orange bars), respectively.

Nucleotide composition near pA sites was analyzed to further validate the reliability of those identified cleavage and polyadenylation sites. In agreement with previous studies [29], Adenine base (A) has higher fraction at -40 to -20 nucleotide (nt) before pA sites while Uridine base (U) peaks downstream of pA sites (Figure 2D). Motif analysis was further performed and classical poly(A) signal (AATAAA at DNA level, PAS) is expected to be the most significant one (Figure 2E). Consistent with the abundance of pA site categories, PAS showed a correlated value of significance (Figure 2E). Furthermore, the nucleotide preference at the site of cleavage is in the order of A>U>C>G for pAs located in different regions of genes (Supplemental Figure S1D). In addition, all pAs identified in this study, except for pAs located in exon, shown very similar features: canonical PAS could be found in the upstream of pAs (Supplemental Figure S1E and S1F) and there was relatively high U-content and lower for C/G-content surrounding the pAs (Supplemental Figure S1F). Such nucleotide preference was also observed in arabidopsis [34] and zebrafish [31]. What is more, frequencies of an additional set of cis elements, which defined by polya_svm [56] and may be involved in the regulation of APA [57], were relatively low in surrounding regions of pAs located in exons (Supplemental Figure S1G). The above results suggest a potential novel mechanism of choosing pAs in exons of eukaryotic genes. Collectively, these data provide strong evidence to support that the pA sites identified by PA-seq are most likely true.

Alternative polyadenylation sites were further investigated for both protein-coding genes and non-coding RNAs (ncRNAs). Our PA-seq analyses manifested that ~42% of all expressed protein-coding genes have two or more PA clusters (Figure 2F, Supplemental Figure S2A-B) and each gene, on average, has 1.66 APA isoforms, which is comparable with another study based on PAS-seq [33] and is significantly higher than the previous estimate based on the entire mouse EST database [55]. In addition, we identified 464 PA clusters covering 345 ncRNAs, and 26% of these ncRNAs have two or more PA clusters and the average number of PA clusters is 1.35 (Figure 2G, Supplemental Figure SC). Some genes with multiple 3′ UTR isoforms may display biased expression to the top isoform. However, this is not generally the case. By excluding the expression percentage of the top isoform, more than 60% of the genes with two or more pAs express the remaining isoforms as a minimum of 10% of the total expression (Figure 2H). What’s more, when excluding the top two isoforms, restricted to genes with three or more pAs, more than 40% of the genes express the remaining isoforms at an appreciable level of 10% of total expression (Figure 2H). Therefore, multiple pAs should be considered at the same time to avoid systematic bias that could be introduced if only two pA sites are considered per gene at a time.

Based on above analyses, top three categories of PA clusters (Known pA, 3′ UTR and extended 3′ UTR) occupy 93% (Figure 2B) of all identified PA cluster numbers and 98% (Figure 2C) of tags inside all identified PA clusters. Therefore, combined tags in these three PA clusters (TPM, or the number of Tags Per Million reads) were used to evaluate the expression level of each gene with APA. To examine whether the PA-seq results reflect relative gene expression level, the same RNA samples were sequenced by strand-specific RNA-seq [58]. In total, we got ~104 million paired-end 101-mer reads, and about 92% of them (~96 millions) were mapped uniquely to mouse genome (see details in Supplementary Table S3). Specifically, we calculated the gene expression from the RNA-seq data with the FPKM (Fragment Per Kilobase of exon model per Million mapped paired reads) value. We only considered genes that expressed in all passages (FPKM ≥ 1 and TPM ≥ 1) for further analysis. PA-seq and RNA-seq profiles were considerably correlated (rho = 0.81, 0.83, 0.79, 0.78 for PD6, PD8, PD10, and PD11, respectively, see Supplemental Figure 3, Supplementary Table S4). Therefore, the read count generated by the PA-seq approach can potentially be used to reflect transcript abundance. Totally, 3,165 genes which have multiple 3′ UTRs caused by APA and with high Spearman’s rank correlation coefficient from PD6 to PD11 between RNA-seq and PA-seq data were used for global APA regulation analyses in cellular senescence.

**Figure3.**
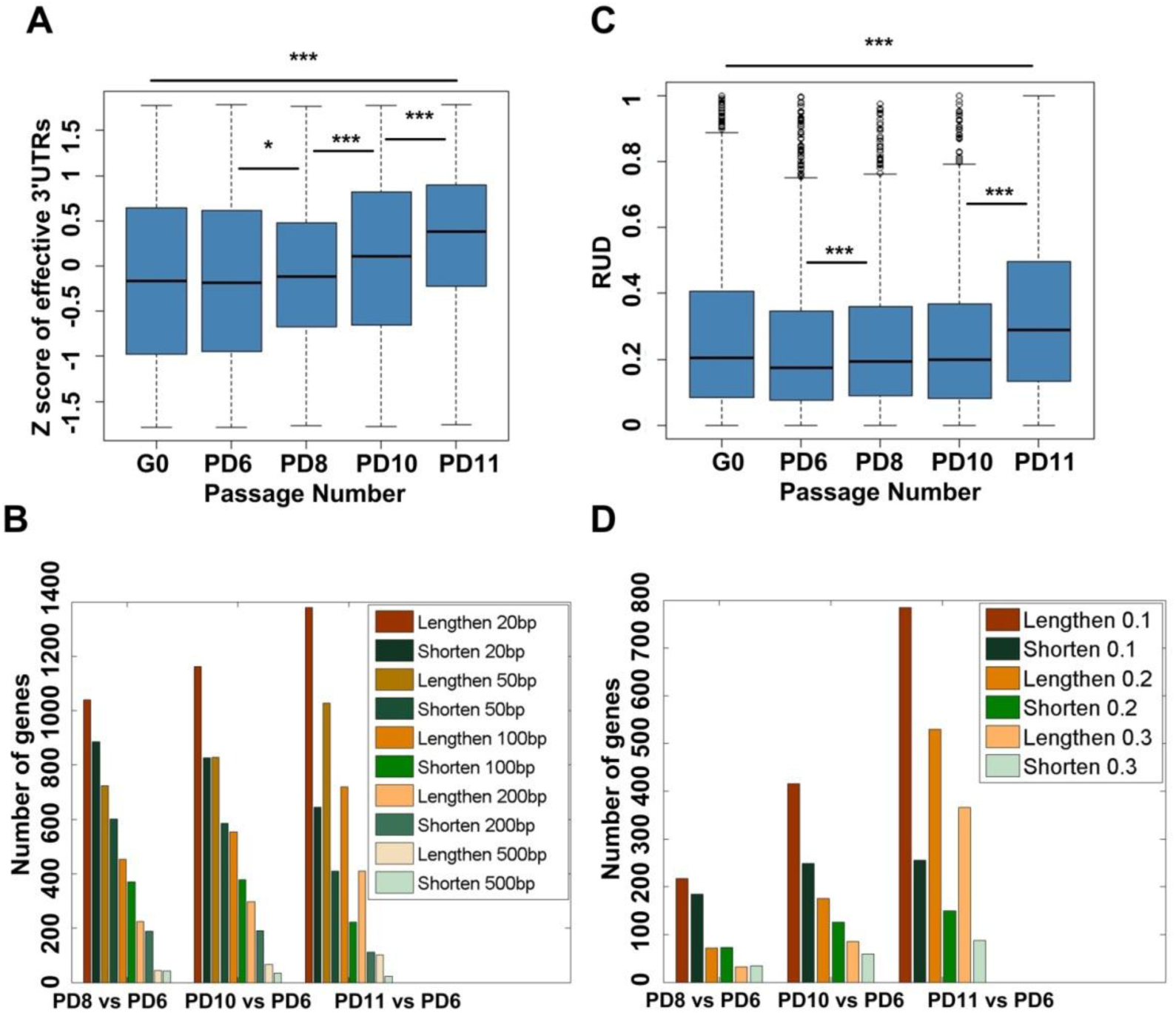
Progressive lengthening of 3′ UTRs for genes with APA regulation during replicative senescence of MEFs. (A) Box plot for Z scores of effective 3′ UTRs across G0, PD6, PD8, PD10 and PD11 of MEFs. (B) Number of genes with lengthened effective 3′ UTRs and number of genes with shortened effective 3′ UTRs by comparing PD11, PD10, PD8 to PD6, respectively, given different thresholds based on PA-seq data. (C) Box plot for RUD scores (relative expression of mRNA isoforms using distal pA sites) across G0, PD6, PD8, PD10 and PD11 of MEFs. (D) Number of genes with higher RUD scores and number of genes with lower RUD scores by comparing PD11, PD10, PD8 to PD6, respectively, given different thresholds based on RNA-seq data. (***) *P* <0.001, (**) *P* <0.01 and (*) *P* <0.05, two tailed Wilcoxon signed rank test. See Materials and methods for more information about Z score of effective 3′ UTRs and RUD score.

### Progressive lengthening of 3′ UTRs for genes with APA during replicative senescence of MEFs

Because our PA-seq data contain experimentally defined pAs as well as their steady-state expression level, it provides us a unique opportunity to interrogate global changes of 3′ UTR length of APA genes among different passages of MEF cells. Based on our published paper [35], effective 3′ UTR length (named Effective_UTR hereafter) was used to estimate the relative usage of pAs inside or downstream of 3′ UTR regions for each gene based on a consolidated gene model by employing both the locations and the tag counts of the PA clusters identified in the 3′ UTR of each transcribed locus (see Methods and Materials) [35]. It should be noted that, this is a considerable improvement over the previous strategies by using microarray or RNA-seq data, which only considered the most proximal and distal PA sites annotated for individual gene loci [52]. We are excited to find that effective 3′ UTR showed a global progressive lengthening trend during the population doubling process of MEF cells (Figure 3A, Supplementary Table S4).

To further evaluate the changes of 3′ UTR length at individual gene level, we compared the effective 3′ UTR length in later passages (PD8, PD10, PD11) with earlier one (PD6) using different cutoff (20bp, 50bp, 100bp, 200bp and 500bp, Figure 3B). There were always more number of genes with longer effective 3′ UTR than with shortened effective 3′ UTR when compared to PD6 at different thresholds (Figure 3B). Noteworthy, the number of genes with lengthened effective 3′ UTR was gradually increased from PD8 to PD11 when compared to PD6, while the number of genes with shortened effective 3′ UTR is gradually decreased from PD8 to PD11 when compared to PD6 (Figure 3B). These results further validated the global lengthening trend during the process of replicative senescence in Figure 3A.

In consistent with the PA-seq data, we also observed the same trend of global 3′ UTR lengthening during replicative senescence in MEFs using the measurement of RUD index based on RNA-seq data by following the analysis pipeline of Zhe et al. [52] (Figure 3C, Supplementary Table S4). The change of RUD index between different passages was applied to evaluate the preference of longer or shorter 3′ UTR for specific gene. In agreement with PA-seq results, more genes have longer 3′ UTR in later passages (PD8, PD10 and PD11) than in earlier passage (PD6) (Figure 3D). In addition, the number of genes with higher RUD index (labeled lengthen in Figure 3D) is continuously elevated from PD8 to PD11 when compared to PD6, while the number of genes with decreased RUD index (labeled shorten in Figure 3D) is gradually reduced from PD8 to PD11 when compared to PD6 (Figure 3D). This independent RUD analysis confirmed the global lengthening of 3′ UTRs during replicative senescence in MEF cells.

To ask whether cell cycle affects effective 3′ UTR length, MEF cells at population doubling time 6 (PD6) were treated with serum starvation to force them enter G0 phase, a reversible cell cycle arrest state. We found the global 3′ UTR pattern of G0 sample is more close to PD6 than to PD11, a more senescent status, by both PA-seq (Figure 3A) and RNA-seq (Figure 3C) data. This result implies that global lengthening of 3′ UTR is more specific to cellular senescence (or irreversible cell cycle arrest) than to reversible cell cycle arrest (G0 phase).

To validate that genes did undergo 3′ UTR lengthening during cellular senescence, we randomly selected 10 genes for quantitative real-time Polymerase Chain Reaction (qRT-PCR) validation (Figure S4-S5, see details about primers information in Supplementary Table S5). Two genes that showed shortened 3′ UTR in PD11 compared to PD6 were included as internal control (Figure S5). All of 10 genes showed an elevated trend of using longer 3′ UTR (Figure S5), with 7 genes have reached a statistical significant level. As a confirmation that qRT-PCR approach did reflect the trend of 3′ UTR length change, all of the two control genes showed a shortened 3′ UTR usage in PD11 compared to PD6 with a statistical significance (Figure S5). The experimental validation supports the observation that senescent cells underwent global 3′ UTR lengthening.

### Longer 3′ UTRs tend to have decreased mRNA abundance during cellular senescence

We next examine the consequence of 3′ UTR lengthening during MEF cellular senescence. Longer 3′ UTR is thought to provide more sites for regulation by microRNA and/or RNA binding protein (RBP) [42, 43, 59], which affects the mRNA abundance and/or translation at post-transcriptional level. In agreement with this hypothesis, a global decrease of gene expression was observed during cellular senescence for genes that have multiple pA sites (Figure 4A). Internal control genes with single pA site did not show such trend (Figure 4B), supporting the noting that global down-regulation of expression level is specific for APA genes. Expression change at individual gene level was also analyzed. To simplify the conclusion, we compared PD11 MEF cell with PD6 cell for both 3′ UTR usage and steady-state expression level. The genes that showed significant lengthened 3′ UTR outnumbered genes containing shortened 3′ UTR four to one (Figure 4C, Figure S6A, Supplementary Table S7). Within 3′ UTR lengthened genes, there are more genes showed a significant downregulation than genes have elevated expression level (*P*<5.6 × 10^−12^, Binomial test, Figure 4C, right half). No such difference was observed for genes with shorter 3′ UTR (Figure 4C, left half). Together, decreased expression at individual gene level supports the observation of global down-regulation of APA genes during cellular senescence.

**Figure4.**
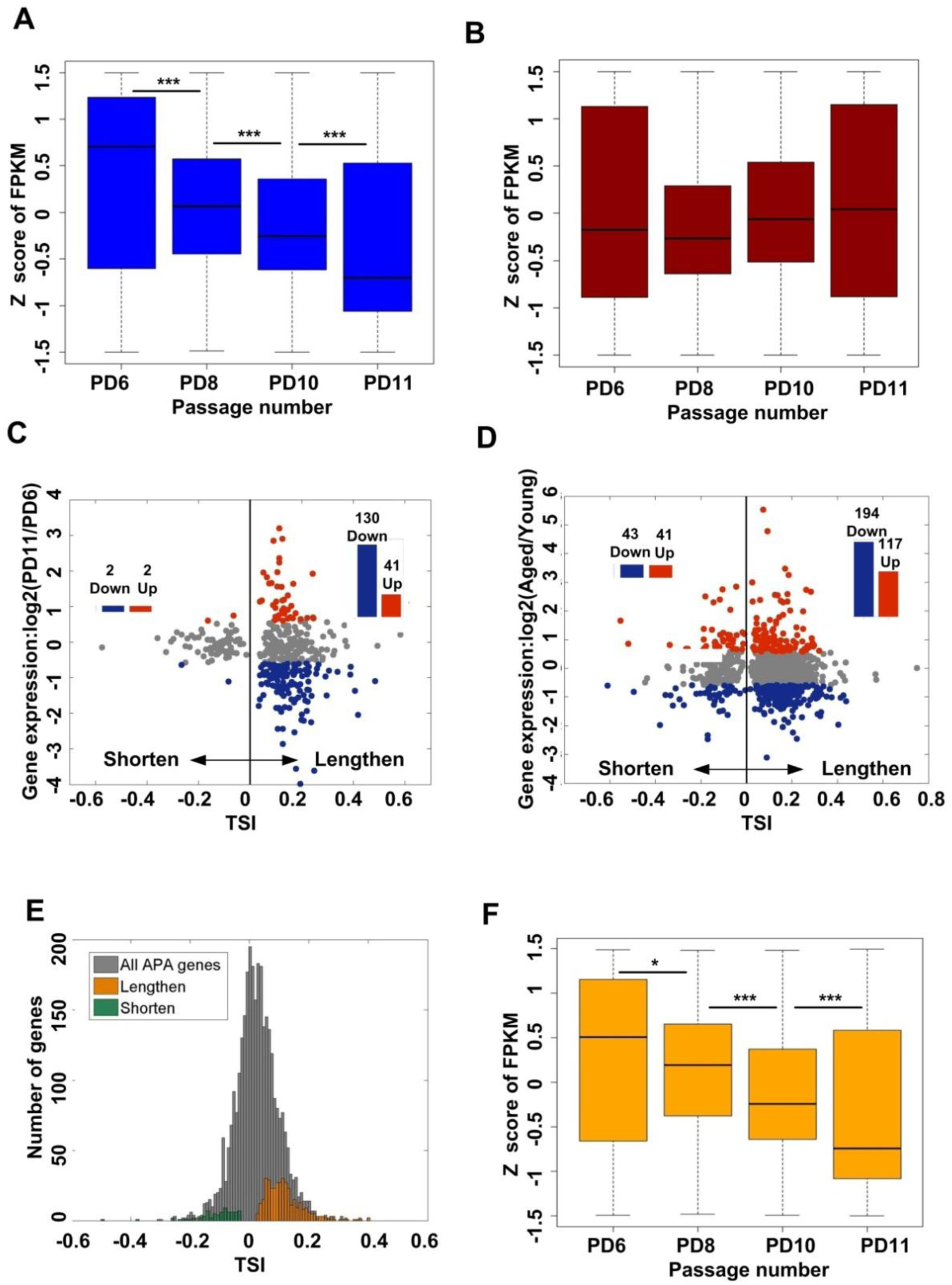
Genes preferred to use distal pAs during cellular senescence tended to have decreased mRNA abundance. (A) Box plot for normalized expression (FPKM, Fragments Per Kilobase of exon model per Million mapped reads) across PD6, PD8, PD10 and PD11 for genes with multiple pAs. (B) Box plot for normalized expression (FPKM, Fragments Per Kilobase of exon model per Million mapped reads) across PD6, PD8, PD10 and PD11 for genes with single pA site. (C) For genes with significantly longer or shorter 3′ UTRs in senescent MEFs (PD11), their fold-change (FC) expression between senescent MEFs and young MEFs (PD6) were plotted against their TSI values. Red and blue represented up- and down-regulated in senescent cells, respectively, which was identified by fold-change more than 1.5 and FPKM>1 in both samples. The genes with significantly longer or shorter 3′ UTRs in senescent MEFs were identified by linear trend test with Benjamini-Hochberg (BH) false-discovery rate at 5%. (D) For genes with significantly longer or shorter 3′ UTRs in VSMCs derived from old rat (2 years old), their fold-change expression between VSMCs of old and young rat are plotted against their TSI values. Red and blue represented up- and down-regulated in senescent cells, respectively, which was identified by fold-change more than 1.5 and FPKM > 1 in both samples. The genes with significantly longer or shorter 3′ UTRs in VSMCs of old rat were identified by linear trend test with Benjamini-Hochberg (BH) false-discovery rate (FDR) at 5%. (E) Distribution of TSI values for all APA regulation genes. Orange bar showed distribution of TSI values for genes using a lengthened 3′ UTR during replicative senescence of MEFs (with a positive Pearson Correlation *r* and FDR<0.05, Linear trend test). Green bar shown distribution of TSI values for genes using a shortened 3′ UTR during replicative senescence of MEFs (with a negative Pearson Correlation *r* and FDR<0.05, linear trend test). Pearson Correlation *r* named as tandem UTR isoform switch index (TSI). (F). Box plot for normalized expression (FPKM) across PD6, PD8, PD10 and PD11 for genes with 3′ UTR lengthening during replicative senescence of MEFs. (***) *P* <0.001, (**) *P* <0.01 and (*) *P* <0.05, two tailed Wilcoxon signed rank test.

To address whether APA regulates gene expression in another aging system, we constructed PA-seq libraries for total RNA from vascular smooth muscle cells (VSMCs) derived from young and aged rats. PA-seq libraries were sequenced and analyzed using the same strategy (see read mapping information in Supplementary Table S6). To our surprise, a very close ratio of longer genes versus shorter genes regarding 3′ UTR length between rat VSMC and mouse MEF cells was observed (Figure S6A, Supplementary Table S7). What’s more, there is a significant overlap of genes showed longer 3′ UTR usage between mouse MEFs and rat VSMCs (Figure S7-S8). Further, the correlation that more genes have down-regulated expression given lengthened 3′ UTR also occurs in rat VSMCs (*P*<1.5 × 10^−5^, Binomial test, Figure 4D). These above results support the notion that APA-medicated 3′ UTR lengthening may play a role in gene expression in multiple cellular senescence systems.

To further ask whether there is a continuous lengthening of 3′ UTR at individual gene level and its consequence on gene expression, linear trend test was applied to identify genes with progressive 3′ UTR length changes in MEFs [31, 60]. There were 375 genes showed a continuous lengthening of 3′ UTR from PD6 to PD11, while only 73 genes had progressively shortened 3′ UTR (Figure 4E, see details in Supplementary Table S7). Interestingly, a similar global expression decrease of these 375 genes correlated with their continuously lengthened 3′ UTR (Figure 4F, Figure S6B). Interestingly, the 73 genes contained shortened 3′ UTR during cellular senescence did not display trend of expression down-regulation (Figure S6C-D). Together, these above results indicated global 3′ UTR lengthening may contribute to decreased mRNA steady-state expression level during cellular senescence.

### Functional Enrichment analysis of genes with longer 3′ UTR during cellular senescence

To address whether 3′ UTR lengthening is related to regulation of the process of cellular senescence, genes with longer 3′ UTR were analyzed for functional enrichment. We first selected genes that showed lengthening in PD11 compared to PD6 since these two passages of MEFs displayed dramatic phenotypic changes (Figure 1F). DAVID (The Database for Annotation, Visualization and Integrated Discovery) from NIAID (National Institute of Allergy and Infectious Diseases, NIH) was applied for the Gene Ontology (GO) and pathway enrichment analyses since this tool is widely used [61]. For the 322 genes that have significant longer 3′ UTR in PD11 compared with PD6 in MEFs, the top enriched GO terms are related to Uniquitin mediated proteolysis, Wnt signaling pathway, Cell cycle, Regulation of actin cytoskeleton and others (Figure 5A, Supplementary Table S8), most of which have connections to cellular senescence ([62–65], see Discussions for details). Genes progressively tend to use distal pAs during replicative senescence of MEFs are also enriched in senescent-related pathways (Figure S9, Supplementary Table S8). Interestingly, most of the top enriched GO terms in rat VSMCs are also related to cellular senescence (Figure 5B, Supplementary Table S8). Strikingly, four of the GO terms (in red for Figure 5A and 5B) are shared between mouse MEFs and rat VSMCs. These shared GO terms or pathways all linked to cellular senescence (see Discussions for details). We further examined whether these two species (mouse and rat) only shared the same pathway but not the same gene or gene itself underwent APA regulation conservetively. Worth noting, six genes in the ubiquitin mediated proteolysis showed longer 3′ UTR in both mouse and rat senescent cells (Figure 5C). There were 7, 2 and 5 genes in the regulation of actin cytoskeleton, cell cycle and wnt signaling pathway, respectively (Figure 5D-F), had the same trend of 3′ UTR lengthening in both mouse and rat senescent cells. These above functional enrichment results supported the idea that an intrinsic link between APA genes and cellular senescence exists.

**Figure5.**
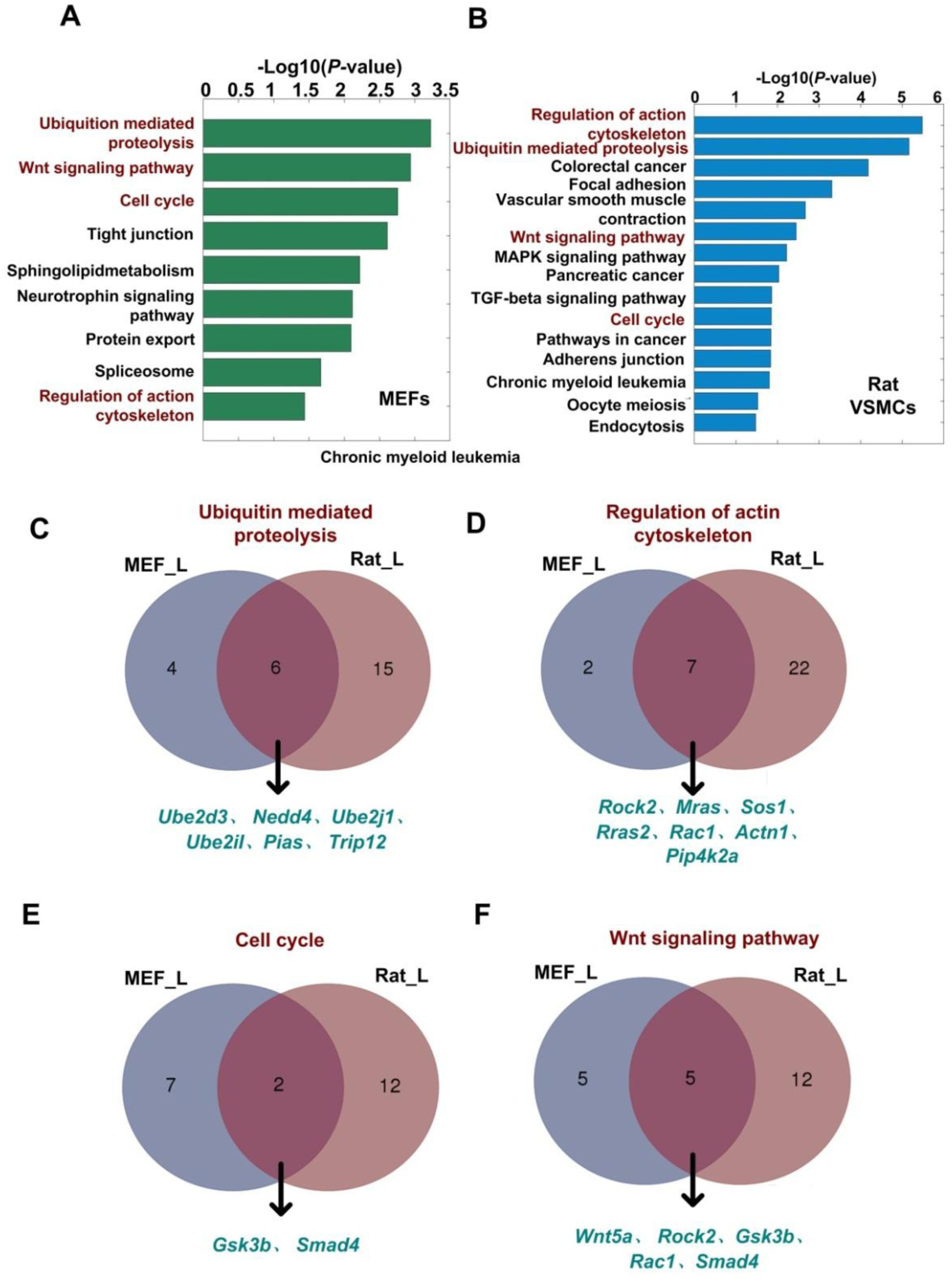
Genes preferred to use distal pAs both in senescent MEFs and VSMCs are enriched in common senescence-related pathways. (A-B) Significantly enriched (*P* <0.01; Fishers’ exact test) pathways of the genes tended to use distal pAs in senescent MEFs and VSMCs derived from old rat, respectively. The common senescence-related pathways between them are marked by red color. (C-F) Venn diagram comparison between mouse and rat for genes involved in ubiquitin mediated proteolysis, regulation of actin cytoskeleton, cell cycle and wnt signaling pathway and preferred to use distal pAs in senescent MEFs and VSMCs of old rat.

### Longer 3′ UTRs introduced more conserved miRNA binding sites and RNA binding protein recognition sequences

To explain why longer 3′ UTR decreases mRNA expression level during cellular senescence, we analyzed the number and density of recognition sites by microRNA (miRNA) and RNA binding protein (RBP) in the alternative 3′ UTR regions between proximal and distal pA sites. In consistent with previous studies, we found that the numbers of conserved miRNA binding sites in long 3′ UTR (determined by distal pA site, named as 3′ UTR_L hereafter) and alternative 3′ UTR (between proximal and distal pA site, named as 3′ UTR_A hereafter) were significantly higher than in short 3′ UTR (determined by proximal pA site, named as 3′ UTR_S hereafter) region (*P*<2.2 × 10^−16^ and *P*<5.6 × 10^−7^, respectively, two side Wilcoxon signed rank test, Figure6A) [43, 59]. Notably, 3′ UTR_L and 3′ UTR_A were also significantly longer than 3′ UTR_S (*P*<2.2 × 10^−16^ and *P*<2.2 × 10^−16^, respectively, two side Wilcoxon signed rank test, Figure6B). Furthermore, the conserved miRNA binding density in 3′ UTR_S was significantly higher than 3′ UTR_L and 3′ UTR_A (*P*<2.2 × 10^−16^ and *P<* 2.4 × 10^−16^, respectively, two side Wilcoxon signed rank test, Figure6C). However, some conserved miRNA binding sites are exclusively located in alternative 3′ UTR (Figure 6D). Interestingly, there was higher density of binding sites in 3′ UTR_L than in 3′ UTR_S for mir-20a-5p, which had been proved involved in cellular senescence of MEFs [66] (*P*<0.03, two side Wilcoxon signed rank test, Figure 6E). In similarity, miR-290a-5p, which acted as a physiological effector of senescence in MEFs[67], also had more higher conserved binding density in 3′ UTR_L and 3′ UTR_A than in 3′ UTR_S (*P*<2.5 × 10^−3^ and *P*<9.4 × 10^−3^, two side Wilcoxon signed rank test, Figure 6F). These above results suggest that more potential miRNA binding sites can be introduced in longer 3′ UTR, although the density of binding can be either high or low in the alternative region.

**Figure6.**
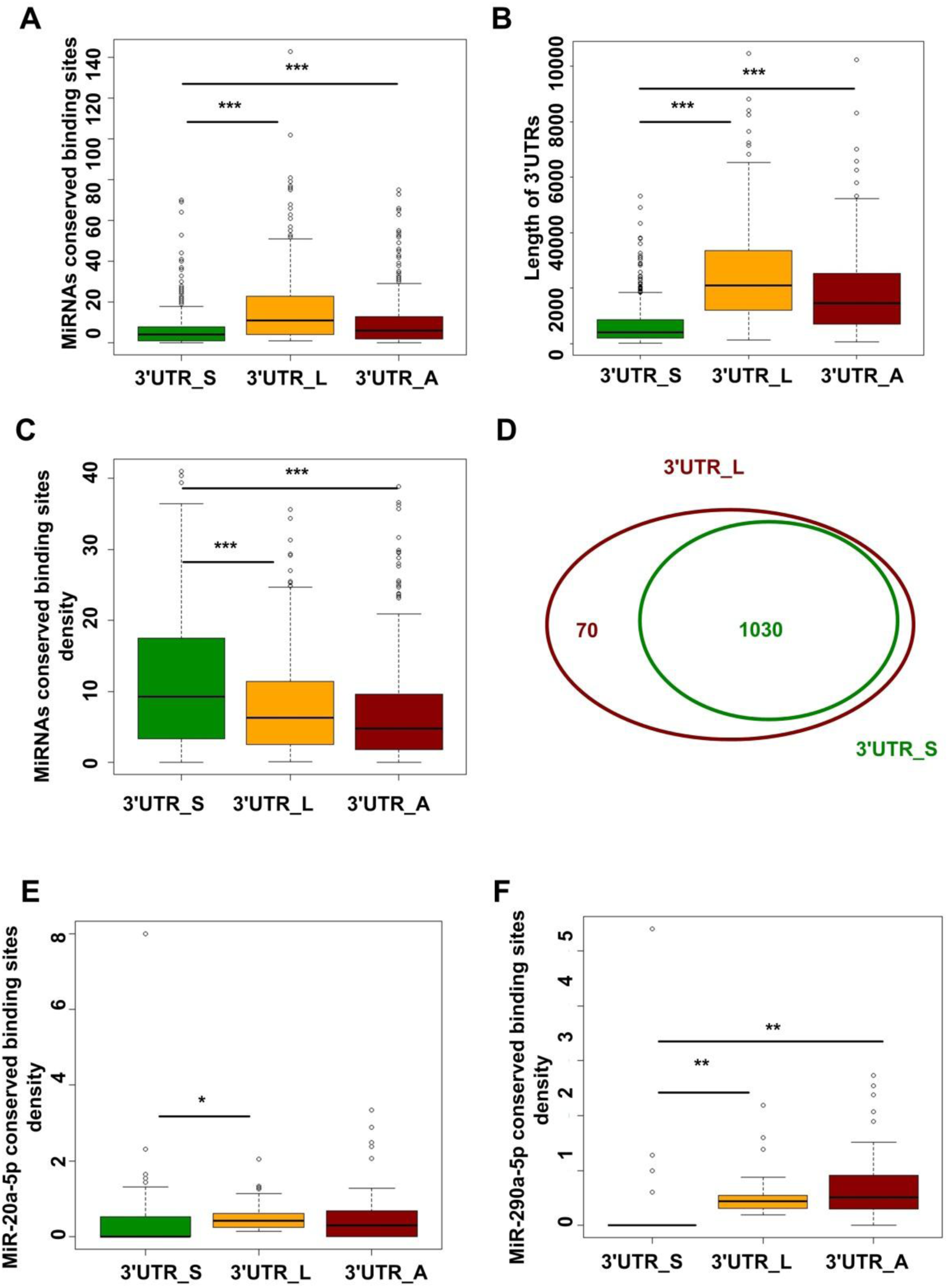
Genes progressively tended to use distal pAs during replicative senescence of MEFs can introduce more conserved miRNA binding sites when use longer 3′ UTRs. (A) Box plot comparison for the number of conserved miRNA binding sites among the shortest 3′ UTRs (named as 3′UTR_S), the longest 3′ UTRs (named as 3′UTR_L) and alternative 3′ UTRs (named as 3′UTR_A) in genes progressively tended to use distal pAs during replicative senescence of MEFs. (B) Box plot for length comparison among 3′ UTR_S, 3′UTR_L and 3′UTR_A in genes progressively tended to use distal pAs during replicative senescence of MEFs. (C) Box plot comparison for density of conserved miRNA binding sites among 3′ UTR_S, 3′UTR_L and 3′UTR_A in genes progressively tended to use distal pAs during replicative senescence of MEFs. (D). Venn diagram comparing conserved miRNA binding sites in the 3′ UTR_S with 3′ UTR_L for genes progressively tended to use distal pAs during replicative senescence of MEFs. (E) Comparing the density of miR-20a-5p potential binding sites in 3′ UTR_S with 3′ UTR_L and 3′ UTR_A for genes progressively tended to use distal pAs during replicative senescence of MEFs. (F) Comparing the density of miR-290a-5p potential binding sites in 3′ UTR_S with 3′ UTR_L and 3′ UTR_A for genes progressively tended to use distal pAs during replicative senescence of MEFs. (***) *P* <0.001, (**) *P* <0.01 and (*) *P* <0.05, two tailed Wilcoxon signed rank test.

The RNA-Binding Protein DataBase (RBPDB) is a collection of experimentally validated RNA-binding sites both *in vitro* and *in vivo,* or manually curated from primary literature [68]. Interestingly, we found that the expression level of about 58% expressed RBPs (307 expressed RBPs with FPKM > 1 in PD6, PD8, PD10 and PD11), annotated by this database, was decreased during replicative senescence of MEFs. RBPmap can predict the binding sites of 94 human/mouse RBPs, 59 among them are also annotated by RBPDB [69]. Using this tool, we found that 30 expressed RBPs had significantly higher conserved binding densities in both 3′ UTR_L and 3′ UTR_A than in 3′ UTR_S (FDR < 0.05, two side Wilcoxon signed rank test, see details in Supplemental Table S9). Specifically, HNRNPA1 [70] and IGF2BP2 [71] can be involved in mRNA translocation, YBX1 [72] can regulate mRNA stability and NOVA1 [72], SNRNP70 [73], HNRNPF [74], HNRNPH2 [75], HNRNPK [76] can regulate APA (Figure 7B-J). Taken together, these above data indicated that longer 3′ UTRs introduced more sites for RNA binding proteins, which may further affect mRNA stability, localization and even contribute to alternative polyadenylation itself.

**Figure7.**
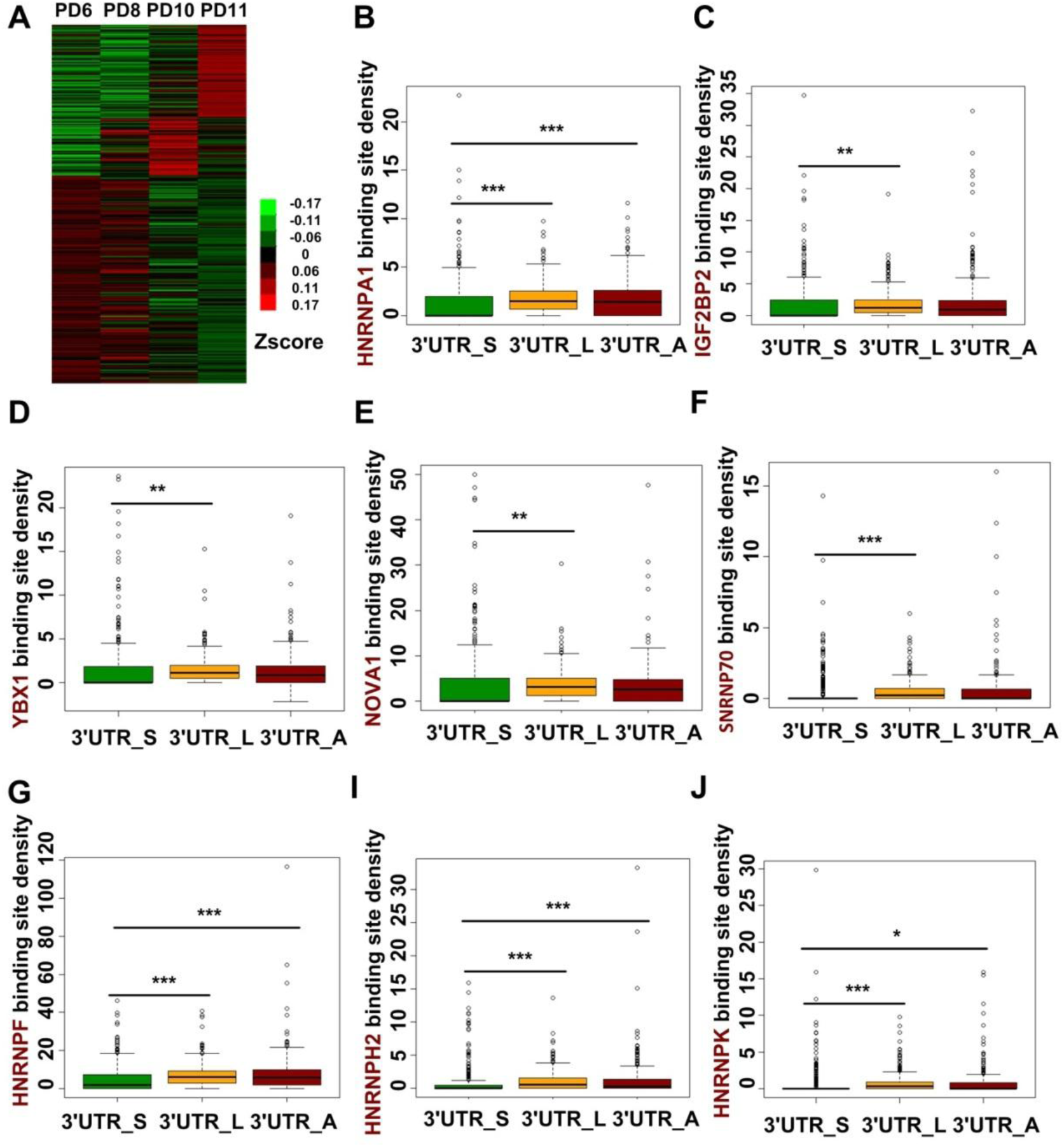
Genes progressively tended to use distal pAs during replicative senescence of MEFs can introduce more conserved RBPs binding sites. (A) Heatmap of RBPs expression during replicative senescence of MEFs. (**B-J**) Comparing the density of HNRNPA1, IGF2BP2, YBX1, NOVA1, SNRNP70, HNRNPF, HNRNPH2 and HNRNPK potential binding sites in the shortest 3′ UTRs (named as 3′ UTR_S) with the longest 3′ UTRs (named as 3′ UTR_L) and alternative 3′ UTRs (named as 3′ UTR_A) for genes progressively tended to use distal pAs during replicative senescence of MEFs. (***) *P* <0.001, (**) *P* <0.01 and (*) *P* <0.05, two tailed Wilcoxon signed rank test.

### Decreased expression of cleavage and polyadenylation related factors may regulate global 3′ UTR lengthening

There are three types of regulators that controlling alternative polyadenylation, including trans-acting RNA binding proteins (RBPs), cis-acting DNA elements and chromatin features near pA sites [38]. Cis elements around proximal or distal pA sites were first analyzed to get insights of the APA regulation during cellular senescence. We compared the sequence conservation around the regions of the proximal and distal pAs of genes that showed progressive lengthening of 3′ UTR (named as LE_proximal and LE_distal, respectively) to other three groups of pA sites, including i) proximal and distal pAs of genes that showed progressive shortening of 3′ UTR (named as SH_proximal and SH_distal, respectively), ii) proximal and distal pAs of genes without significant changes of pAs usage preference (named as NC_proximal and NC_distal, respectively) and iii) pA regions of genes only contain one pA site (named as Single pA). Interestingly, polyadenylation signal (PAS) upstream LE_distal pAs was more conserved than that of LE_proximal pAs, implying that reduced usage of shorter 3′ UTR during cellular senescence may be mediated by weaker proximal PAS (Figure 8A). Supporting this notion, the percentage of ‘AAUAAA’, the strongest PAS, in LS_distal pA sites is the highest among all these seven types of pA sites (Figure 8B). In addition, the downstream sequence conservation scores of proximal pA sites (including LE_proximal, SH_proximal and NC_proximal) are obviously higher than that of distal pA sites (including LE_distal, SH_distal and NC_distal), this may attribute to lower frequencies of strong PAS in proximal pAs than distal pAs, therefore needed to provide more binding sites for RBPs to help they be used (Figure 8B), and also may attribute to 3′ UTR is more conserved than intergenic regions, given downstream sequence of proximal pA site is still in the alternative 3′ UTR region (Figure 8A). Interestingly, among three types of proximal pA sites and three types of distal pAs, downstream sequence conservation score of pAs with significantly usage shift (LE_proximal, SH_proximal, LE_distal, SH_distal) are more conserved than pAs without significantly usage shift, this may also attribute to more binding sites for RBPs needed to be supply to help these pAs with significantly usage shift to be used (Figure 8A). Furthermore, among three types of proximal pA sites, downstream sequence conservation score of LE_proximal is higher than SH_proximal and NC_proximal (Figure 8A), raising the possibility that genes showed lengthening of 3′ UTR during cellular senescence were regulated by providing more conserved binding sequence between proximal and distal pA sites.

**Figure8.**
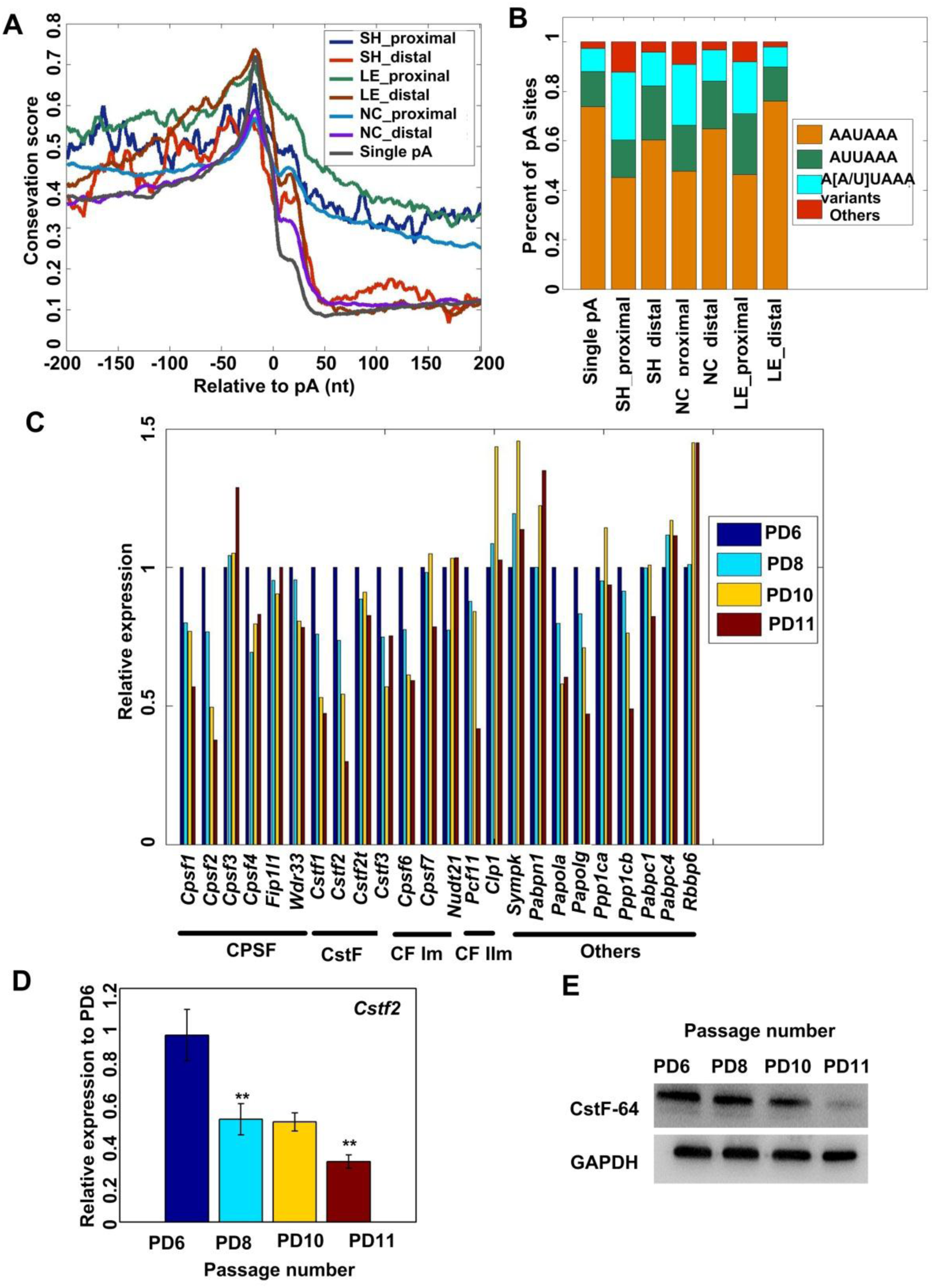
Potential mechanisms for APA regulation during replicative senescence of MEFs. (A) Comparison of conservation score in the 400nt region surrounding the proximal pAs and distal pAs of genes showed continuous 3′ UTR shortening during replicative senescence of MEFs (named as SH_prxomal and SH_distal, respectively), the proximal pAs and distal pAs of genes showed continuous 3′ UTR lengthening during replicative senescence of MEFs (named as LE_prxomal and LE_distal, respectively), the proximal pAs and distal pAs of genes without significant pAs usage preference during replicative senescence of MEFs (named as NC_prxomal and NC_distal, respectively) and pAs of genes with only single pA site. (B) Distribution of PAS sequences in the -40 to -1 nt region for different types of pAs. (C). Gene expression of known polyadenylation factors during replicative senescence of MEFs based on RNA-seq data. (D) Expression levels of *Cstf2* in PD6, PD8, PD10 and PD11 passages of MEFs were detected by qRT-PCR. (E) Detection protein levels of CstF-64 (gene symbol is *Cstf2*) in PD6, PD8, PD10 and PD11 passages of MEFs by Western Blot.

We next examined the possibility that RBPs control APA during cellular senescence. While the core factors in the 3′ processing machinery play essential roles in the cleavage and polyadenylation (C/P) reaction, regulation of their expression level had also been implicated in APA regulation [27]. Therefore, the gene expression of 22 important poly(A) trans-factors [81] were analyzed based on the MEF replicative senescence RNA-seq data (PD6, PD8, PD10, PD11). Consistent with the overall expression trend of RBPs, we observed global down regulation of most poly(A) trans-acting factors during replicative senescence (Figure 8C), in which more genes tend to use longer 3′ UTR. This result is in line with a recently published study, in which global upregulation of most poly(A) factors is associated with more 3′-UTR-shortening events in five tumor types compared with normal tissues [48]. Another survey of the constitutive components of the polyadenylation machinery and other candidates from the literature revealed several trans-acting factors that were significantly upregulated in the cancer cell lines compared with non-transforming cells [44]. Specifically, consistent with the idea that cellular senescence can be a defense system for cells on the way to becoming cancerous [8], we noticed mRNA level of Cstf2, which was the first core C/P factor that can regulate APA [77], was down-regulated during MEF cellular senescence (Figure 8C) and up-regulated in cancer cells [44, 48]. Furthermore, Yao et al. showed knockdown of CstF64 (coded by *Cstf2* gene) leaded to more genes favored distal pA sites, which contributed to lengthening of 3′ UTR in human HeLa cells [78]. We thus validated the mRNA expression and protein production of *Ctsf2* gene during MEF cell senescence. qRT-PCR (Figure 8D) and Western Blot (Figure 8E) confirmed that *Cstf2* did show a reduced trend of mRNA level and protein expression, which may contribute to the 3′ UTR lengthening during cellular senescence. Notably, Li et al. found that *Pcf11* could enhance usage of proximal pAs and *Pabpn1* could promote usage of distal pAs by using siRNA knockdown coupled with deep sequencing [79]. Interestingly, the expression of *Pcf11* was down-regulated and *Pabpn1* was up-regulated during replicative senescence of MEFs, consistent with the global 3′ UTR lengthening pattern. Taken together, we provided evidence that changed expression of cleavage and polyadenylation related factors such as CstF64, Pcf11 and PABPN1, may regulate global 3′ UTR lengthening during MEF cellular senescence, although the speculation deserves further experimental validation in MEF cells.

## DISCUSSION

An increasing number of reports demonstrate the prevalence of global APA regulation in a wide range of biological processes and pathological diseases [26, 39, 48]. For instance, more genes favor distal pA site usage, leading to global 3′ UTR lengthening in cellular differentiation and development [42]. However, upon cellular proliferation, neuron activation and malignant transformation, more genes tend to favor proximal pA site usage, resulting in global 3′ UTR shortening [43, 44, 46, 80]. Cellular senescence is regarded as a cancer prevention mechanism and a contributing factor for individual ageing[8, 9]. Nevertheless, whether global APA regulation exists in this important process is completely unclear. We thus applied our PA-seq method to profile global APA events by using mouse embryonic fibroblast (MEF) replicative senescence as a model system [11–14]. If grown under standard tissue culture conditions, MEFs proliferate limited number of population doublings (PD) and eventually undergo replicative cellular senescence[12, 14]. DNA damage, losses of centrosomal integrity and accumulation of mutations due to oxidative stress may also contribute to the entry of MEFs intro senescence in culture [14]. DNA damage response (DDR) is further proved to be activated when MEFs undergo replicative cellular senescence [12]. However, the genome-wide molecular consequence of such signal is unclear.

Interestingly, global 3′ UTR lengthening during MEF replicative senescence was observed. What’s more, we found the expression levels of genes with multiple pAs showed a decreased steady-state expression during cellular senescence. Furthermore, genes both preferred to use distal pAs in senescent MFEs and showed progressive lengthening of 3′ UTR during MEF cellular senescence have a decreased trend of steady-state mRNA expression level (Figure 4). In contrast, this gene expression decreasing pattern in senescent cells cannot be observed for genes either with single pA site or tended to use proximal pAs (Figure 4B and Figure S6D). The observation that genes used proximal pA sites in senescent cells did not show a trend of increased expression is consistent with the result that genes preferred longer 3′ UTR in cancer cells did not have a decreased trend of expression[48]. Since global lengthening of 3′ UTR in senescent cells and global shortening of 3′ UTR in cancer cells were observed, these above results support the notion that APA-mediated 3′ UTR lengthening serves as a posttranscriptional regulation layer in MEF cellular senescence.

To address whether this phenomenon of 3′ UTR lengthening in cellular senescence is specific for mouse or can extend to other species, we sequenced and analyzed PA-seq data derived from old and young rat’s aortic vascular smooth muscle cells (VSMCs). Intriguingly, there are also more genes prefer distal pAs usage in elder rat than in younger rat (Figure 4D), suggesting global 3′ UTR lengthening in older cells is evolutionary conserved. What is more, genes preferred distal pAs also tended to display a trend of decreased gene expression. Most strikingly, shared pathways (Figure 5A,B) and genes (Figure 5C,D,E,F) preferred to use distal pAs both in senescent MEF cells and in rat VSMCs derived from older rat were also be found. Consistent with the above results, Xia et al. observed broad shortening of 3′ UTRs across seven TCGA (The Cancer Genome Atlas) tumor types and that genes with shorter 3′ UTRs in tumors were prone to be up-regulated [48]. Furthermore, many genes were found to have opposite pAs usage preference between senescent cells and cancer cells (Supplemental Figure S10). On the one hand, this was consistent with the previous knowledge that senescence response was a potent tumor suppressive mechanism [8]. On the other hand, this also implied that APA regulation may play a vital role in deciding cell destiny, although definitely worth further intensive investigation.

Excitingly, genes with longer 3′ UTR in senescent MEF cells and rat cells derived from older rat’s vascular smooth muscle are all enriched in common pathways such as ubiquitin mediated proteolysis[62, 81, 82], regulation of actin cytoskeleton[65, 83], cell cycle[64] and wnt signaling pathway[63, 84], which was proved to be involved in cellular senescence and cell cycle arrest (Figure 5 and Supplemental Figure S9). Interestingly, the genes which were preferred to use proximal pAs and with increased expression levels in cancer cells were also enriched in ubiquitin mediated proteolysis and regulation of actin cytoskeleton pathways[48]. This also implied that the APA regulation of genes involve in these two pathways may play some important role in cell fate decision, such as senescence or tumorous.

Proteostasis is maintained by the proteostasis network, which comprises pathways that control protein synthesis, folding, trafficking, aggregation, disaggregation, and degradation [85]. Notably, the identification of shared “ubiquitin mediated proteolysis” pathways between mouse and rat old cells is of interest as these machineries play a vital role in protein homeostasis and their functionality declines with age [85–87], while perturbations of the ubiquitin-proteasome pathway have been involved in pathogenesis of various neurodegenerative disease including Alzheimer’s disease (AD) and (Huntington disease) HD [88, 89]. Specifically, NEDD4, an E3 ubiquitin ligase regulated by APA both in replicative senescence of MEFs and senescent rat VSMCs (Figure 5C), was shown to delay cellular senescence by degrading PPARγ and eventually upregulating SIRT1[90]. Interestingly, NEDD4 was also be proved to be a potential proto-oncogene that negatively regulated a conserved component of the IIS (insulin/insulin-like growth factor-1-signaling) cascade, the phosphatase PTEN via ubiquitination, a paradigm analogous to that of Madm2 and p53[91]. In addition, Cul1, which belong to a Skp1-Cul1-F-Box E3 ligase complex, also involved in regulating the lifespan of *Caenorhabditis elegans* [92]. Interestingly, a Cul1 complex has also been shown to inhibit the activity of FOXO proteins by catalyzing their degradation [93–95]. Cul1 also can promote cell-cycle progression at the G1-S phase transition through regulating the abundance of cyclin E [96]. Furthermore, the full-length 3′ UTR of *Cull* gene, which was proved to use distal pAs in senescent MEFs by qRT-PCR (Figure S11A-B), has significantly lower stability (*P*= 3.5 × 10^−3^, *P*= 1.5 × 10^−3^ and *P*= 2.2 × 10^−3^ for 4, 8, 12 hours after actinomycin D treatment, respectively, two-tailed t-test, Supplementary Figure S11C). To assay how different 3′ UTR isoforms of *Cul1* influence protein expression, shorter, full-length and full-length with depletion of the sequence containing the proximal PAS (named as 3′ UTR_S, 3′ UTR_L and 3′ UTR_M, respectively) were fused to a dual-luciferase reporter assay system. Both 3′ UTR_L and 3′ UTR_M yielded lower luciferase activity than the construct containing only the common (or shorter) 3′ UTR regions in NIH3T3 mouse fibroblast cell line (*P*<0.04 and *P*<0.02, respectively, two-tailed t-test, Supplementary Figure S11D). These above data further support the idea that APA may be involved in cellular senescence via affecting the mRNA stability and protein translation efficiency of genes in senescence-related pathways.

Gourlay et al. demonstrated that increasing actin dynamic, either by a specific actin allele or by deletion of a gene encoding the actin-bundling protein Scp1p, can increase lifespan of yeast by over 65%, which may due to lower ROS level was produced [97]. In addition, CDK5 activation was upregulated in senescent cells, which further accelerated actin polymerization and induced senescence and the senescent shape change by reducing GTPase Rac1 activity [98]. Florian et al. also found that an unexpected shift from canonical to non-conical Wnt signaling in mice due to elevated expression of Wnt5a in aged haematopoietic stem cells (HSCs) appeared to induce stem cell ageing [99]. Interestingly, we found both Rac1 and Wnt5a tended to use distal pAs both in replicative senescence of MEFs and senescent rat VSMCs. In conclusion, our presented studies have revealed that APA regulation may be involved in several signaling pathways which are, likely, implicated in the molecular phenotypes of cellular senescence. However, whether these alterations reflect responses to the progression of ageing or causal mechanisms should await further experimentation.

In agreement with previous studies, we found that alternative 3′ UTR regions between proximal and distal pA sites contained higher numbers of conserved miRNA seed matches than common (i.e. shorter) 3′ UTR regions (Figure 6A) [43]. The alternative 3′ UTRs also contain 70 unique miRNA binding sites that do not appear in the common region (Figure 6D), providing additional cis elements for posttranscriptional regulation. Interestingly, the average length of alternative 3′ UTR is larger than common 3′ UTR (Figure 6B), leading to the density of potential seed matches of miRNAs in alternative 3′ UTR is lower than that of common 3′ UTR (Figure 6C). It is worth noting that the overall number is more important than the density of miRNA binding sites in posttranscriptional regulation. Excitingly, a few senescence-associated miRNAs even have both larger number and higher density of potential miRNA binding sites in alternative 3′ UTRs. As examples, miR-20a-5p and miR-290–5p, both of which have higher conserved miRNA binding density in alternative 3′ UTR regions (Figure 6E,F), have been proved to induce senescence in MEFs [66, 67]. In addition, miR-93 and miR-494, which also expressed in MEFs based on Nidadavolu et al.’s data and contained higher seed binding site densities in long 3′ UTRs based on our data, also involved in senescence of MFEs [100–102]. Besides miRNAs, many RBPs, which may regulate the stability or translocations of mRNA [70–72], have higher densities in full length (i.e., long) and alternative 3′ UTRs than in short 3′ UTR regions (Figure 7). Therefore, using distal pAs in senescent MEFs provide multiple opportunities for miRNAs and RBPs binding with 3′ UTR regions of mRNAs, which may regulate mRNA stability, translation and translocation.

In consistent with previous investigations, we found that the distal pAs which preferred to be used in senescent MEFs tend to have more conserved and more strong poly(A) signal (PAS) [26, 44]. Notably, the correlation of using distal pAs with generally down-regulated of core factors in C/P reactions is in agreement with the observation that up-regulation of core factors is associated with proximal pAs preference in cancer cells [44, 48]. In general, we speculate that a combination effect of the expression level changed RBPs eventually determined the global 3′ UTR lengthening, although some of the RBPs may favor proximal pA site usage or have context dependent roles for different genes [103–105]. As a considerable number of gene’s APA can be affected by certain RBPs, it is reasonable that some of these RBPs can have observable phenotypic changes upon knockdown. Taken together, all these above results raise an intriguing model for the regulation role of APA in cellular senescence. Upstream signals down-regulate most of the RBPs, including core RBPs for cleavage and polyadenylation, some of which have significant roles in controlling the APA of genes, thus leads to a global lengthening of 3′ UTR; Longer 3′ UTRs trend to contain more miRNA binding sites and/or more cis-elements that affect RNA stability and/or translation efficiency, eventually affect the protein level of genes that related to cellular senescence. However, more experimental validations are warranted to fully prove such hypothesis.

Notably, there are two major categories of cellular senescence including replicative senescence and stress-induced premature senescence (SIPS) [6]. Here we proved that MEFs replicative senescence underwent global 3′ UTR lengthening. VSMCs derived from old rat are likely a combination of replicative senescence and varieties of stress-induced senescence. Although we discovered more genes use longer 3′ UTR, whether SIPS itself will induce global 3′ UTR lengthening remains elusive. Thus more cellular senescent models are required to fully understand the prevalence and functional relevance of 3′ UTR lengthening, which serves as an important posttranscriptional gene regulation mechanism. Importantly, PA-seq and RNA-seq only can manifest the steady-state level of mRNAs, nascent transcript sequencing such as GRO-seq[106] and Bru-seq[107], may tell us whether the genes tended to use distal pAs during transcription in senescent cells. In addition, in this study, we only predicted the potential binding sites for miRNAs and RBPs in 3′ UTR regions. miRNA expression data and also CLIP-seq [108] or PAR-CLIP seq [109] data for RBPs can help us further investigate how miRNAs and RBPs shape the mRNA fate after transcription. Furthermore, TAIL-seq, which allows us to measure poly(A) tail length at the genomic scale, is also a potent tool that can be used to dissect dynamic regulation of mRNA turnover and translational control [110]. Recently, Lars et al. developed BAT-seq to analyze single-cell isoform data, and found that single cells from the same state differed in the choice of APA isoforms [111]. In this study, we only investigated the pA usages of genes from a cell population, whether a more senescent cell is more prone to use distal pA can be further investigated by using the same or comparable method.

In conclusion, these above results provide evidences to support the intrinsic link between APA and senescence in multiple species, although functional validation and underlying mechanism deserves further investigation.

### Materials and Methods

#### Cell isolation and cultivation

For replicative senescence, primary MEFs isolated from embryos of 12.5–14 days’ pregnant C57BL/6 mice according to the method described previously [62] were cultured in DEME (GIBICO, Grand Island, NY, USA), supplemented with 10% fetal bovine serum (FBS, GIBCO, Grand Island, NY, USA) and 1% Penicillin/Streptomycin (GIBICO, Grand Island, NY, USA), at 37 °C in a humidified incubator with 5% CO2. To develop replicative senescence, confluent MEFs were evenly transferred into new dishes and the cells were cultured until get confluent again to generate one population doubling (PD). MEFs were maintained in a 100-mm dish, and in this study, confluent status of cells means almost confluent culture condition without contact inhibition. The number (n) of PD was calculated using the equation (*n* = log_2_ (*Ne*/*Ns*), where Ne and Ns are the numbers of cells at the end of cell culture and those seeded at the start of one passage, respectively. The numbers of population doublings (PD, times) and total culturing period were monitored one time every three days until MEFs completely stopped proliferating.

#### Cell cycle analysis by FACS

MEFs were grown in 100-mm dishes and harvested with 1 ml of 0.25% trypsin when reached 70–80% confluence. And then cells were washed twice with ice-cold 1× PBS, fixed with 70% ethanol for 2 hours in 4 °C and washed again with the same PBS before penetrated with 1% Triton X-100 plus RNase A (40 U/ml) and stained with propidium iodide (50 p,g/ml) on shaking platform at 37 °C. As staining finished, DNA contents were measured on a FACS Calibur HTS (Becton Dickinson). The percentage of diploid cells in G1, S and G2/M phases were analyzed by Summit version 5.0 software (Dako, Campbellfield, Australia).

#### SA-β-gal staining

SA-β-gal activity, as a classic biomarker of senescent cells, was also monitored at the same time for all the growth stages of MEFs. Culture medium was removed from 12-well plates and cells were washed once with 1 ml of 1× PBS and fixed with 0.5 ml of fixative solution for 10–15 minutes at room temperature. While the cells were in the fixative solution, the staining solution mix (staining solution, staining supplement and 20 mg/ml X-gal in DMSO) was prepared according to the manufacturer’s instructions (Biovision, Cat. #K320–250, USA). As washed twice with 1 ml of 1× PBS, cells were incubated with the aforementioned mix at 37 °C overnight or 12 hours. Then the cells were observed under an inverted microscope (Leica, Germany).

#### qRT-PCR and Western Blot

MEFs were harvested at 70–80% confluence. Total RNA and protein were isolated with TRIzol Reagent according to the provided manual (Invitrogen, USA). For qRT-PCR analysis, RNA was reverse-transcribed into cDNA using random primers, and mRNA levels of *Mki67, p16* and *Cstf2* were measured by qPCR (LightCycler, Roche, Swiss). See Supplementary Table S10 for primer information.

For western blot, protein was resolved in 1% SDS, subjected to SDS-PAGE, transferred to nitrocellulose membrane and incubated with primary antibodies of interest (p16, 1:500–1000, sc-1207, Santa Cruz; Cstf2, 1:1000, ab64942, abcam; GAPDH, 1:2000–3000, sc-32233, Santa Cruz) overnight at 4 °C. In the second day, the blots was washed three times in PBST (1× PBS + 0.1% Tween-20) and incubated with diluted secondary antibodies (IRDye800CW Goat anti-Mouse (1:10000, Cat.926–32210, LI-COR) for 1–2 hours at room temperature. And then the protein blots were washed three times with PBST before visualized under odyssey infrared imaging system (Odessey, LI-COR).

#### Validation of pAs usage preference by qRT-PCR

RNAs were reverse transcribed with oligo(dT), followed by PCR with two pairs of primers (proximal and distal) targeting different regions of the cDNAs. Specificity, the region targeted by the proximal pair is common to both APA isoforms and the region by the distal pair is unique to the longer isoform. qPCR signals from the proximal and distal pairs of primers are compared to indicate the relative expression of the two isoforms. See Supplementary Table S3 for primer information.

#### mRNA stability

NIH3T3 Cell lines were treated with actinomycin D (5 μg/ml), and after 0, 4, 8, 12 and 24 hours total RNA was isolated and analyzed by qRT-PCR. Primer sequence can be found in Supplementary Table S5.

#### Luciferase Assays

psiCHECK-2 Vector (Promega, Cat. no. C8021) provides a quantitative and rapid approach for measuring the effect of 3′ UTR on protein translation efficiency. Renilla luciferase is used as a primary reporter gene, and the 3′ UTR of interest can be cloned downstream of the Renilla luciferase translational stop codon. Measurement of Renilla luciferase activity is a convenient indicator of 3′ UTR influencing on protein translation efficiency while the transcription level is unchanged. This firefly reporter cassette has been specifically designed to be an intraplasmid transfection normalization reporter. Thus, a short 3′ UTR, a long 3′ UTR and a long 3′ UTR with a mutation of proximal polyadenylation signal can be compared.

NIH3T3 cells were seeded into a 24-well plate at a density of 15,000 cells/well, 3 kinds of Mouse Cull 3′ UTR were separately subcloned into the psiCHECK-2 Vector using the XhoI and PmeI restriction enzyme sites. After an overnight incubation, the cells were treated with a transfection mixture consisting of 50 μl of serum-free medium, 1 μl of Lipofectin reagent (life technologies, Cat. no. E2431), 0.5 μg of psiCHECK-2 Vector:short; 0.5 μg of psiCHECK-2 Vector:long; 0.5 μg of psiCHECK-2 Vector:mutant each 4 well. After a 20-minute incubation, 100 μl of serum-containing medium was added to the wells. 24 hours post transfection Renilla and firefly luciferase activities were measured using the Dual-Luciferase Reporter 1000 Assay System. The results reflect the influence of 3′ UTR on protein translation efficiency indirectly.

#### PA-seq and RNA-seq library construction

The PA-seq libraries were constructed by following the protocol published before by Ni et al.[35]. The dUTP-based strand-specific RNA-seq libraries were constructed by following the protocol of Parkhomchuk et al [58]. Both library types were sequenced by Illumina HiSeq2000 platform in a paired-end 2×101 bp manner.

#### Paired-end mapping and mRNA abundances estimation

Paired-end reads from PA-seq were subjected to strand corrected according to ‘TTT’ at the beginning of the reads as previously described [35]. FastQC software was used for quality control of PA-seq and RNA-seq data (http://www.bioinformatics.bbsrc.ac.uk/projects/fastqc/). Processed raw data were then aligned to mouse genome (version mm9) or rat genome (version rn5) using Tophat2 [112]. Reads were mapped uniquely using the “-g 1” option. Bowtie was used to get a good approximation for -r and --mate-std-dev parameters, which were needed for Tophat2 [113]. Combined counts of all PA clusters mapped to annotated PA, 3′ UTR and extended 3′ UTR regions (TPM, or the number of Tags Per Million reads) were used to evaluate the expression level of these genes with APA regulation. RNA-seq reads were also aligned to the genome in the similar way and Cufflinks was used to assemble transcripts, estimate the mRNA abundances by following the instruction published before [114].

#### Peak calling

After mapping, the information of cleavage and polyadenylation sites (pAs) was extracted. All aligned reads were pooled together so that a unified peak-calling scheme can be applied. F-seq [53] was used for peak calling of pAs by default parameters setting expect that set the feature length as 30. We resized the PA clusters to the shortest distance that contained 95% of the reads according to our previous publication [35]. To filter internal priming, we removed PA clusters with 15 ‘A’ in the 20 nucleotides region downstream of peak mode or with continuous 6 ‘A’ in the downstream of peak mode. Peaks with fewer than 20 supporting reads were removed. RefSeq annotation was used for the genomic location analysis. We classified the genome into eight groups and mapped the pA peaks to these groups using the following hierarchy: PA > 3′ UTR > Extended 3′ UTR > Exon > Intron > 5′ UTR > TSS > Promoter. Especially, PA defined as annotated pA site plus upstream and downstream 10 bp. Extened_3′ UTR denoted downstream 5 kb of 3′ UTR by following the previous study [35]. Note that the peaks mapped to extended 3′ UTR region should land in intergenic region and not overlapped with any known transcripts. TSS (Transcription Start Site) represented exact transcriptional start site plus upstream and downstream 10 bp. Promoter defined as upstream 250 bp of TSS, which should also not overlap with any known transcripts. Next, we also wanted to exclude the situation where a weak and seldom used APA signal generates an event that passes the expression threshold simply because the gene itself is highly expressed by following the previous study [115]. We therefore kept cleavage events that accounted for ≥ 10% of the total reads mapped in the corresponding genes in ≥ 1 samples. If an event was lower expressed but was expressed by > 5% of the total reads mapped in the corresponding genes in the majority of samples (≥ 80%), we also kept the pA site.

#### Comparison of PA-seq data with PolyA_DB

PolyA_DB2 was downloaded from Bin Tian’s website (http://exon.umdnj.edu/polya_db)[54]. They retrieved all cDNA/EST sequences listed in human, mouse, rat, chicken and zebrafish UniGene databases from NCBI (July and August 2005 versions), and then aligned them with genome sequences downloaded from the UCSC Genome Bioinformatics Site (http://genome.ucsc.edu) using BLAT[54]. The genome version be used here is mm5 for mouse, therefore we used the online tool LiftOver (http://genome.ucsc.edu/cgi-bin/hgLiftOver), which can convert genome coordinates and genome annotation files between different assemblies, to change the pAs to the version of mm9. Due to the imprecise nature of cleavage by CPSF-73 or fuzzy definition of the pAs by their surrounding *cis* elements, the cleavages may occur at different positions within a small sequence window [116]. Therefore, if the pAs identified by PolyA_DB2 located in the upstream 10 bp or downstream 10 bp region of the pAs identified by our data, we take them as the same pAs.

#### Motif analysis

MEME motif enrichment analysis was used to identified the enriched motifs in the upstream (-40 nt) of our identified pAs, with a motif length was set to 6 which is equal to the length of PAS [117]. In addition, frequencies of matches of 15 motif identified by Hu et al. [57] was defined by polya_svm, only highly similar matches of motifs were considered (> 75th percentile of all possible positive scores) [56].

#### Define the effective 3′ UTR length and RUD score

For each expressed genes, the effective 3′ UTR length and RUD score based on the PA-seq and RNA-seq data were calculated, respectively. We used effective 3′ UTR length to analyze the PA-seq data following our previous study [35]. Specificity, we used Entrez gene ID to cluster the RefSeq transcripts IDs. Overlapping genes on the same strand were removed in order to avoid misassignment of pAs to genes. We also restricted our attention to genes whose annotated transcripts shared the same stop codon and also without junctions in the 3′ UTR regions. Based on the consolidated gene model above, the pipeline considers both the corresponding 3′ UTR length and the tag counts of the pA peaks identified in the tandem 3′ UTR regions, including pA peaks located in known pAs, 3′ UTR and also extended 3′ UTR regions, of each transcribed locus to compute “effective 3′ UTR length” [35]. For genes located in the plus strand, the 3′ UTR length for each pA was calculated by the right boundary of the pA peak minus the stop codon. While, for genes located in the minus strand, the 3′ UTR length for each pA was calculated by the stop codon minus the left boundary of the pA peak.

The calculation of RUD (the score for relative expression of isoforms using distal PolyA sites) based on RNA-seq was also following a previous study by Ji et al [52]. Only uniquely mapped and properly paired reads were used for the calculation of RUD. Properly paired reads are those with two pairing reads mapped to different strands of the same chromosome. RUD was based on the ratio of read density in aUTR (alternative UTR) to that in cUTR (constitutive UTR). The regions upstream and downstream of a proximal pA were named as cUTR and aUTR, respectively. The proximal pAs were defined by PA-seq data. Therefore, a high RUD value manifests higher abundance of long 3′ UTR isoforms resulting from usage of promoter-distal pAs. In theory, the range of RUD should be from 0 to 1. However, the RUD may be more than 1 caused by either low or biased coverage of RNA-seq reads in 3′ UTR regions, therefore the genes with RUD more than 1 would be discard also. A standard score (Z score) of Effective 3′ UTR length and RUD for each gene across different passages were calculated for normalization. In addition, a cutoff of both TPM = 1 (Tags of Per Million mapped reads) and FPKM = 1 (Fragments Per Kilobase of exon model per Million mapped reads) for PA-seq and RNA-seq in each passage respectively, was used to select expressed genes.

#### Comparison of APA profiles between different passage of MEF cells

The 3′ UTR switching for each gene among the four passages was detected by measuring the linear trend alternative to independence similarly descried as previously reported [31, 60]. To test a gene with totally *J* tandem pAs across *I* passages, we performed the test as follows: 1) calculate the tandem 3′ UTR length for each of the *J* tandem pA; 2) put the number of reads for each tandem pA at each passage to an *I* × *J* contingency table: take tandem pAs as columns (from the site with shortest 3′ UTR to that with longest); take the *I* stages by time series as rows; 3) if the total number of reads in the *I* × *J* table was less than 30, this gene was discarded for the test; otherwise 4) let the lengths of tandem 3′ UTR denote the scores for the columns; let the row number denote the row score; 5) calculate Pearson correlation *r* (named as tandem 3′ UTR isoform switch index, TSI, following the paper of Li et al [31]) by the following formula:

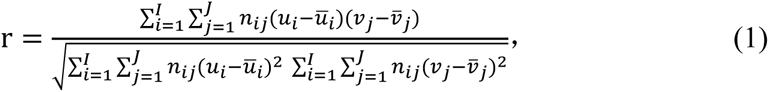

where 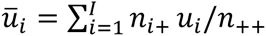 and 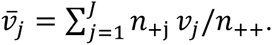.

6) calculate the test statistic *M*^2^ = (*n*_++_ − 1)*r*^2^; For large samples, this statistic is approximately chi-squared with df = 1 and *P* value can be obtained. The genes with significant *P* value corresponding to a false discovery rate (FDR) cutoff of 5% (Benjamini-Hochberg), estimated by Benjamini-Hochberge method with R software, were considered as significantly different among different passages. In specificity, a FDR value less than or equal to 0.05 with positive TSI implies a lengthening 3′ UTR across the different passages of cellular senescence; FDR value less than or equal to 0.05 with negative TSI implies a shortening 3′ UTR.

#### Pathways enrichment analysis

DAVID (Database for Annotation, Visualization and Integrated Discovery) [61] was used for the pathways enrichment analysis and KEGG (Kyoto Encyclopedia of Genes and Genomes) database was chose.

#### Prediction conserved binding sites of miRNA and RBP in 3′ UTRs

The sequences of all mature miRNAs of mouse were downloaded from the miRBase database (Release 21). 8mer, 7mer-m8 and 7mer-1A targets denote conventional miRNA seed matches [118]. Conserved target sites were identified using multiple genome alignments obtained from UCSC Genome Browser (mm9, 30-way alignment) requiring a perfectly aligned match in mouse, rat, human and dog, same as the paper of Sandberg et al[43]. Highly conserved RBP binding sites were predicted by RBPmap online tools.

#### Cluster analysis of RBPs

Clustering of genes based on their FPKMs was done by using the Cluster software (http://rana.lbl.gov/EisenSoftware.htm). We first log transformed FPKM of each gene in each passage, centered them by the mean and then normalized each gene. K-means clustering approach with euclidean distance was used to classify the various types of gene profiles. The cluster result was then visualized by Treeview software (http://taxonomy.zoology.gla.ac.uk/rod/treeview.html).

#### Conservation analysis surrounding pAs

Conservation scores in the 400nt surrounding different types of pAs were calculated using phastCons tracks from 30 vertebrates and was downloaded from UCSC genome browser (http://hgdownload.soe.ucsc.edu/goldenPath/mm9/phastCons30way/)[119].

##### Data Access

Both PA-seq and RNA-seq raw data can be found at the NCBI Sequence Read Archive (SRA) with submission number SRP065821.

## ACKNOWLEDGEMENTS

We thank Dr. Jun Zhu for critical reading and insightful comments of the manuscript. This work was supported by National Key Basic Research Program of China (973 program: 2013CB530700, 2015CB943000), National Science Foundation of China (31271348, 31471192), Shanghai Pujiang Talent Plan (13PJ1400700), the Thousand Talents Plan and Research and Innovation Project of Shanghai Municipal Education Commission (14ZZ007).We thank Genergy Biotech (Shanghai) Co., Ltd. for the deep sequencing service.

## SUPPLEMENTAL DATA

**Figure S1, related to Figure 2.**
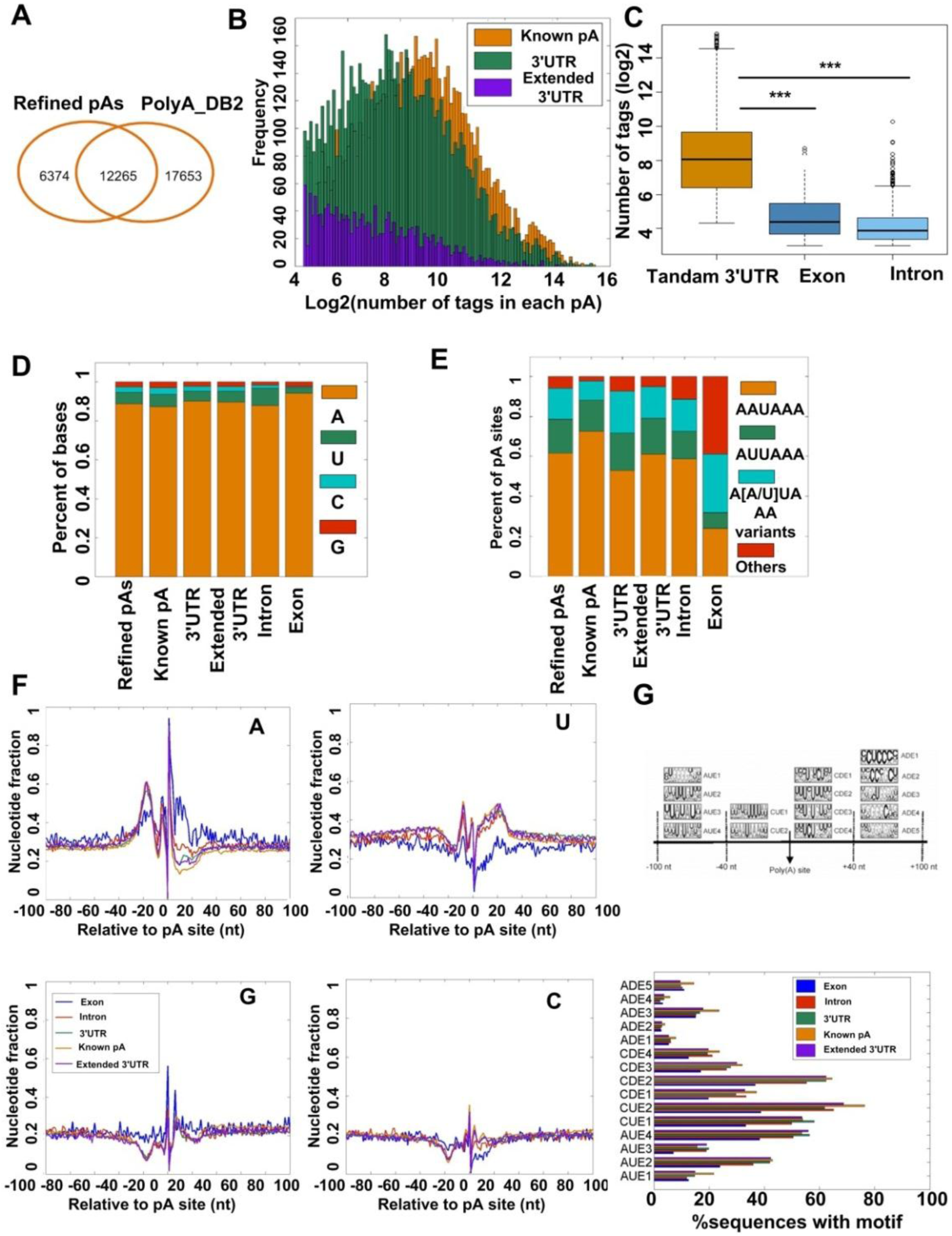
Base composition comparison of different types of pAs. (A) Venn diagram comparing pAs in the PolyA_DB2 database with those identified in this study. (B) Distribution of the number of tags from different types of tandem pAs. (C) Box plot for comparing the number of pA tags in tandem 3′ UTRs with pA tags located in exon and intron. (D) Comparison of base composition of the 3′ base next to pAs from different genomic locations. (E) Distribution of PAS sequences in the -40 to -1 nt region of pAs from different genomic locations. (F) Base compositions in the 200nt surrounding pAs from different genomic locations. (G) Frequencies of matches of different motifs defined by Hu et al. in the 200nt surrounding pAs from different genomic locations was predicted by polya_svm.[1, 2]. Only highly similar matches of motifs were considered (>75th percentile of all possible positive scores). (***) *P* <0.001, (**) *P* <0.01 and (*) *P* <0.05, two Mann-Whitney U test.

**Figure S2.**
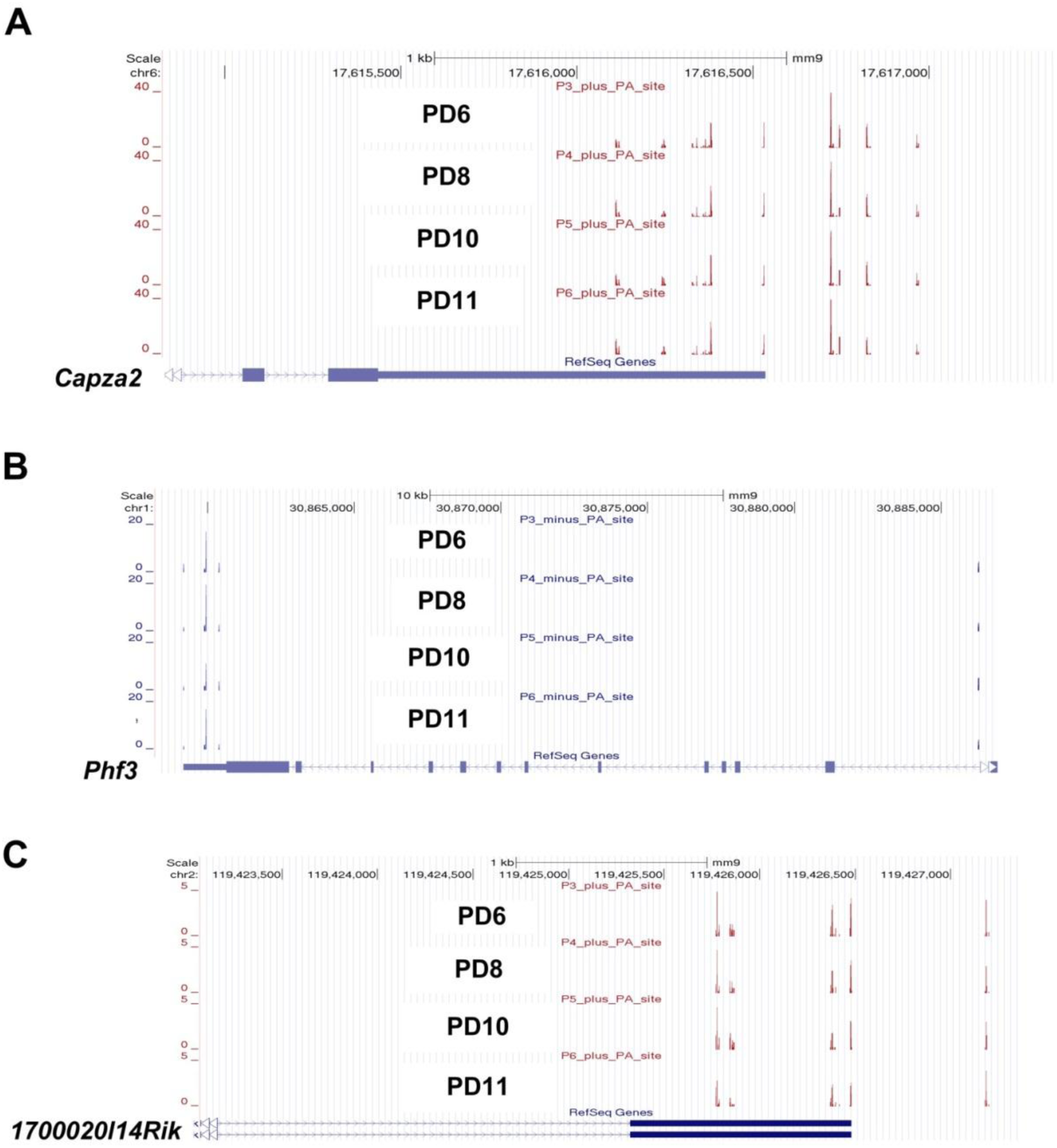
Examples of dynamic APA. (A-C) PA-seq tracks of three genes (*Capza2, 1700020114Rik* and *Phf3*) with multiple pAs.

**Figure S3.**
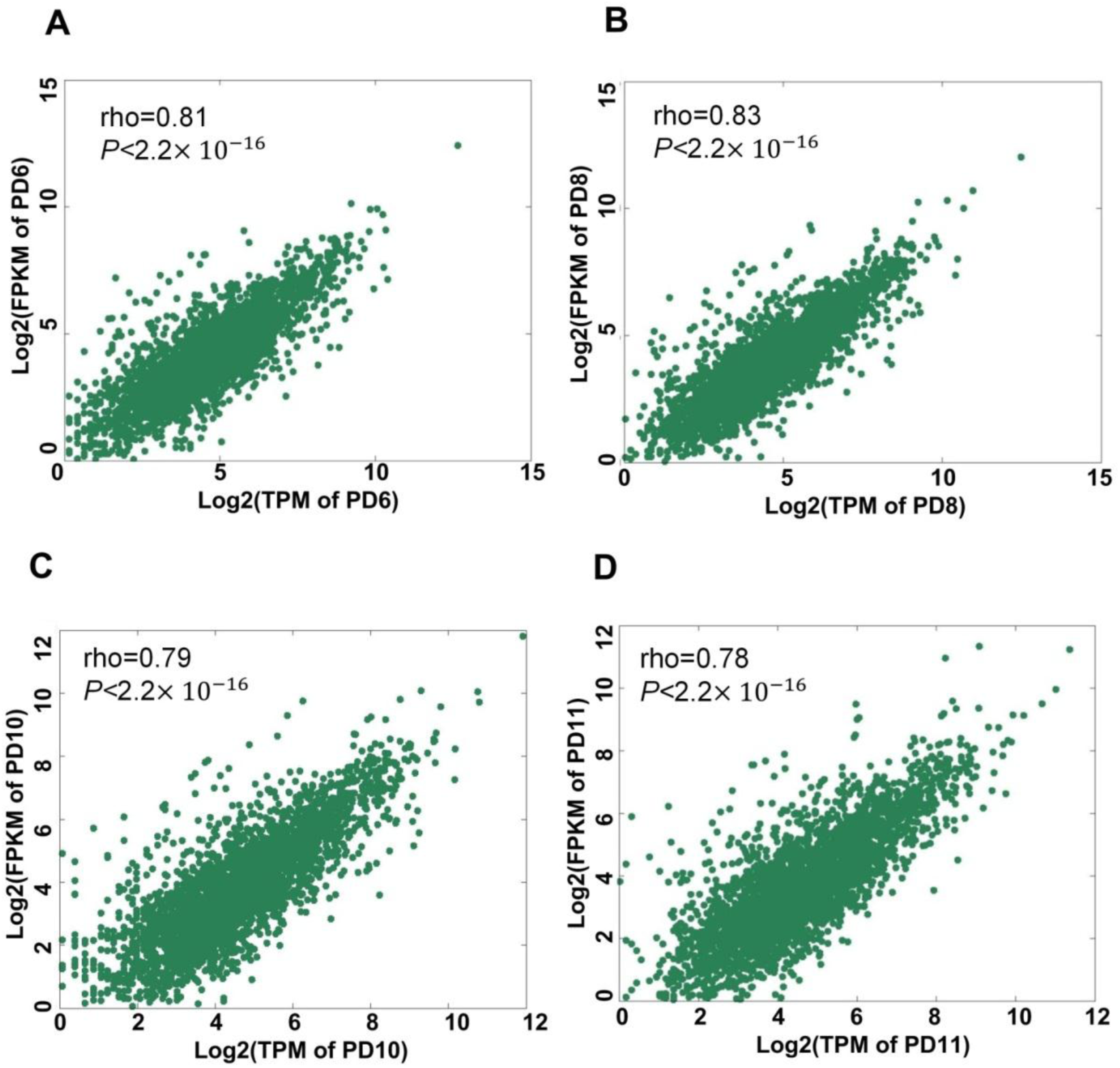
Correlation of expression measured by PA-seq and RNA-seq for genes with APA regulation. (A-D) Correlation of expression measured by PA-seq and RNA-seq for genes with APA regulation in PD6, PD8, PD10 and PD11, respectively. Rho means Spearman’s rank correlation coefficient.

**Figure S4.**
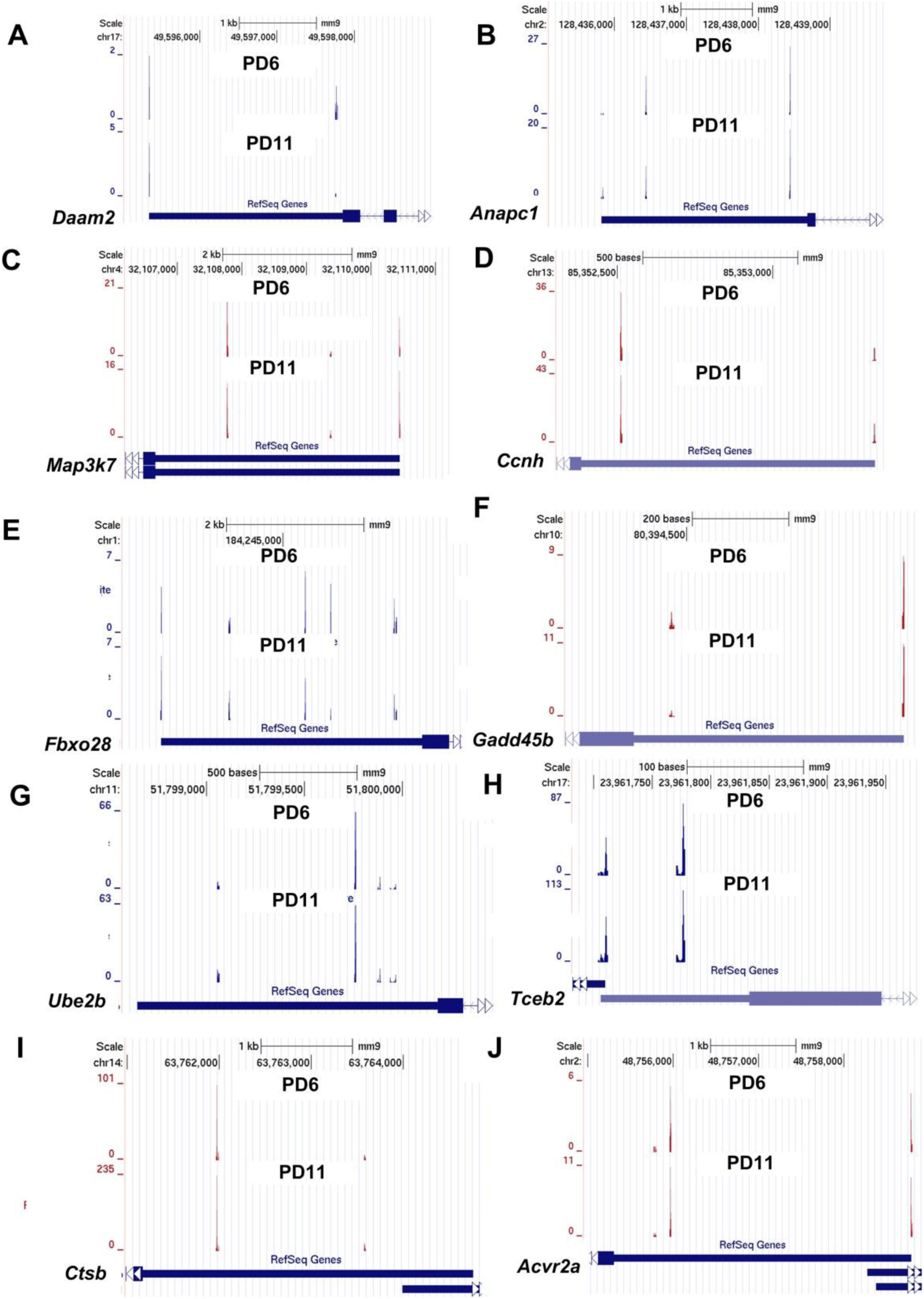
PA-seq track of genes randomly chosen to be validated if they tended to use distal pA sites significantly in senescent MEFs by qRT-PCR. (A-J) PA-seq tracks of *Daam2, Anapc1, Map3k7, Ccnh, Fbxo28, Gadd45b, Ube2b, Tceb2, Ctsb* and *Acvr2a* in senescent and young MEFs, respectively.

**Figure S5.**
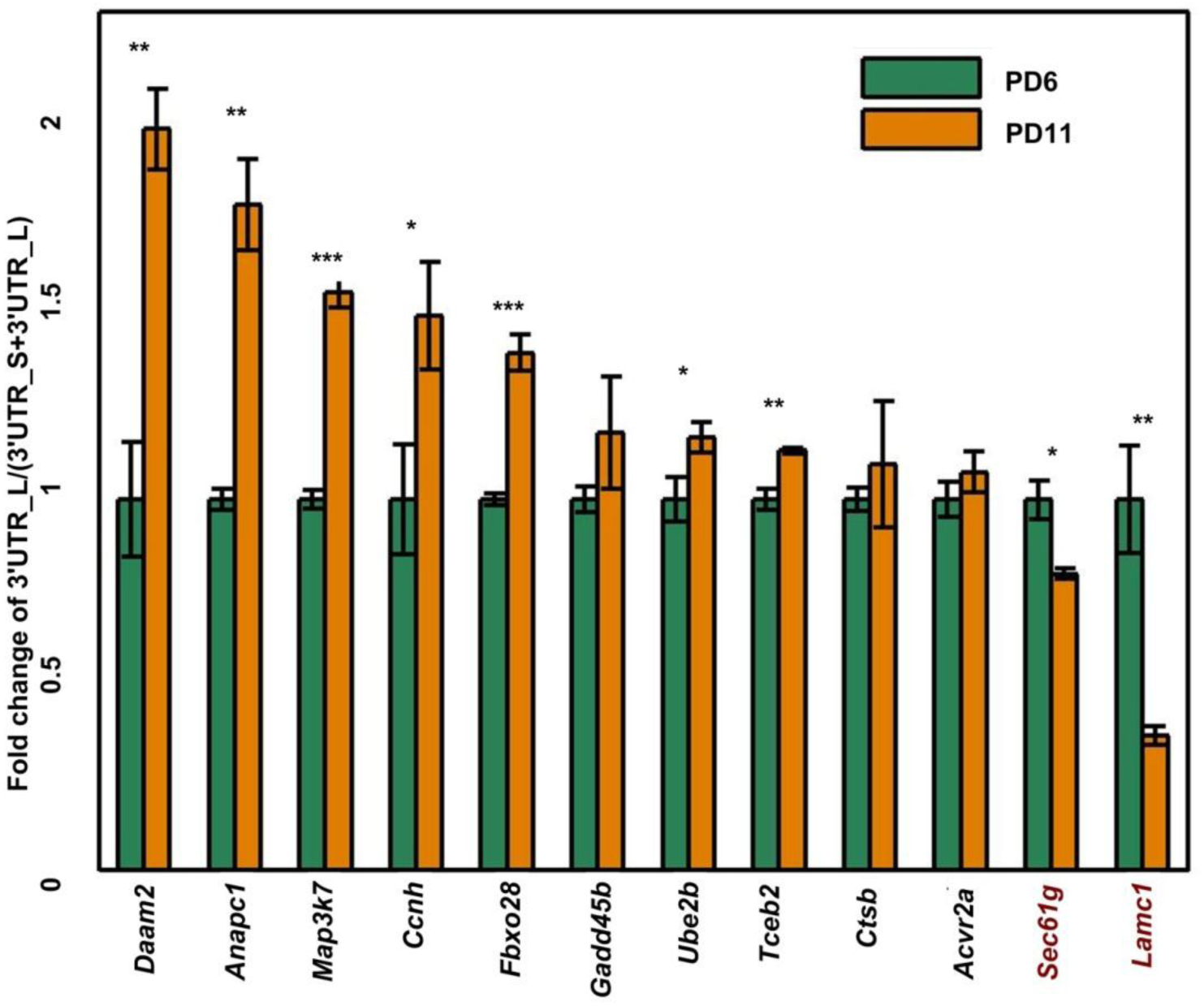
Validation results of genes tended to use distal pAs in senescent MFEs by qRT-PCR. Two genes (*Sec61g* and *Lamc1*) preferred proximal pAs in senescent cells served as negative controls. (***) *P* <0.001, (**) *P* <0.01 and (*) *P* <0.05, two tailed Student’s t-test.

**Figure S6.**
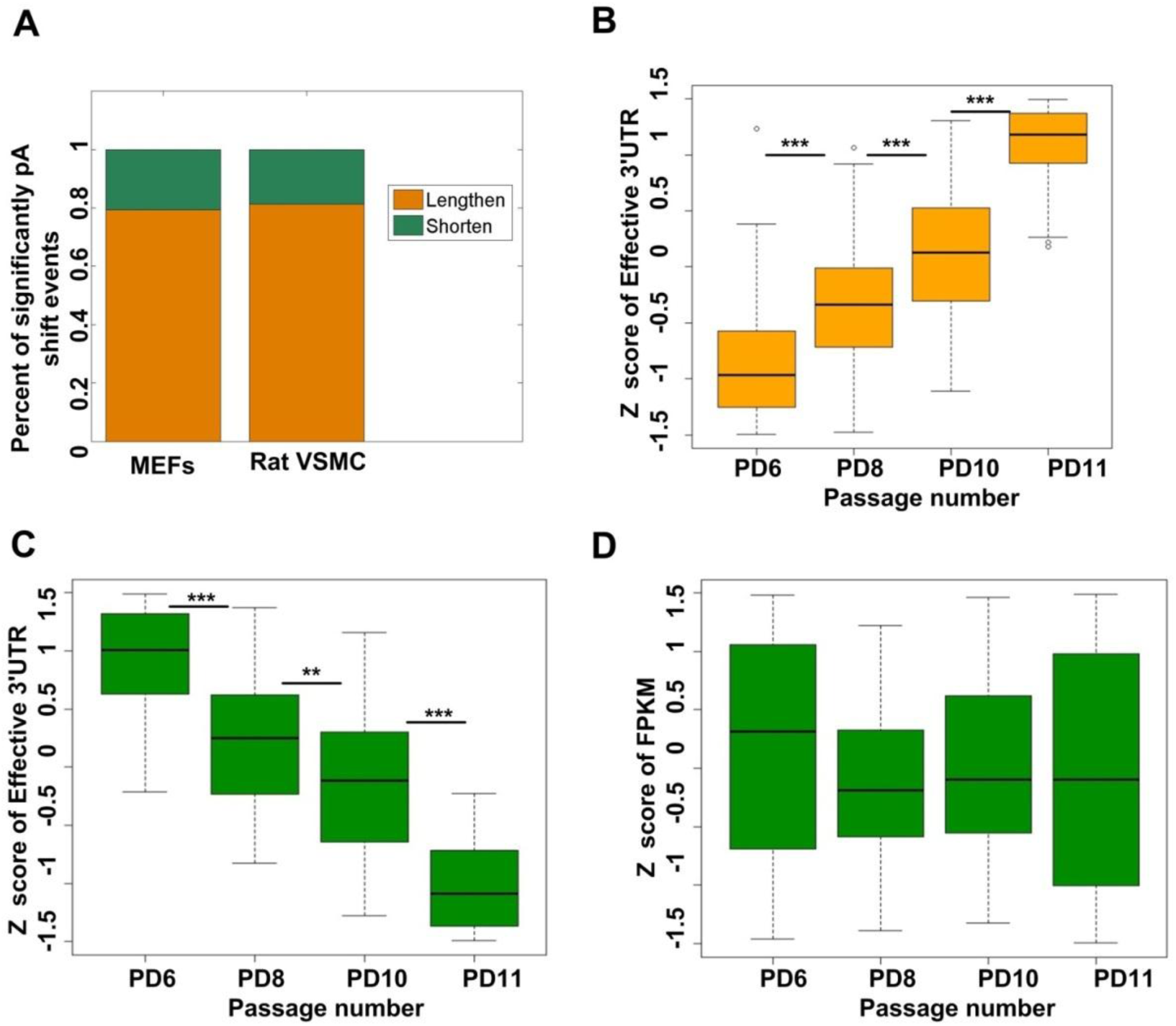
related to Figure 4. Genes tended to use proximal pAs did not have decreased mRNA abundance. (A) Fraction of genes significantly tended to use distal pAs (Lengthen) and proximal pAs (Shorten) when comparing senescent MEFs (PD11) with early passage of MEFs (PD6) and also fraction of genes significantly tended to use distal pAs (Lengthen) and proximal pAs (Shorten) when comparing VSMC (Vascular smooth muscle cells) from old rat (2 years old) with young rat (2 weeks old). (B) Comparing distribution of Z score of effective 3′ UTRs across PD6, PD8, PD10 and PD11 for genes with lengthened 3′ UTRs during replicative senescence of MEFs. (C) Comparing distribution of Z score of effective 3′ UTRs length across PD6, PD8, PD10 and PD11 for genes with shortened 3′ UTRs during replicative senescence of MEFs. (D). Comparing distribution of Z score of expression (FPKM) across PD6, PD8, PD10 and PD11 for genes with shortened 3′ UTRs during replicative senescence of MEFs. (***) *P* <0.001, (**) *P* <0.01 and (*) *P* <0.05, two tailed Wilcoxon signed rank test.

**Figure S7.**
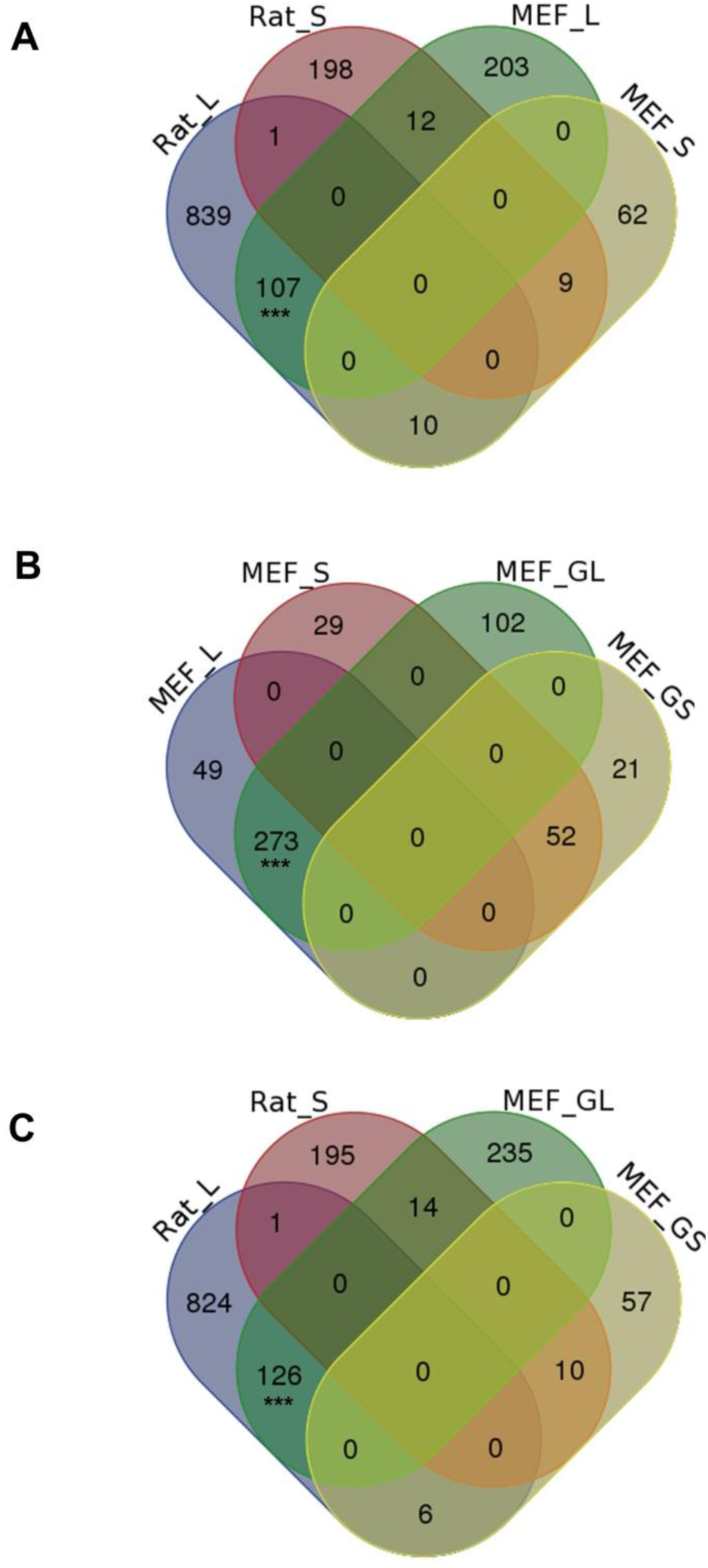
Comparison among genes significantly tended to use distal pAs and proximal pAs in senescent MEFs and VSMCs of old rat. (A) Venn diagram for genes preferring distal (named Rat_L) and proximal (named Rat_S) pAs when comparing VSMCs of old rat with young rat (named as Rat_L and Rat_S, respectively) and genes preferring distal (MEF_L) and proximal (MEF_S) pAs when comparing senescent MEFs (PD11) with Young MEFs (PD6). (B) Venn diagram comparison among MEF_L, MEF_S and genes gradually preferred to use distal (MEF_GL) and proximal (MEF_GS) pAs during replicative senescence of MEFs. (C) Venn diagram comparison among Rat_L, Rat_S, MEF_GL and MEF_GS. MEF_L, MEF_S, Rat_L, Rat_S, MEF_GL and MEF_GS were identified by linear trend test with Benjamini-Hochberg (BH) false-discovery rate (FDR) at 5%. (***) *P* <0.001, (**) *P* <0.01 and (*) *P* <0.05, Fishers’ exact test.

**Figure S8.**
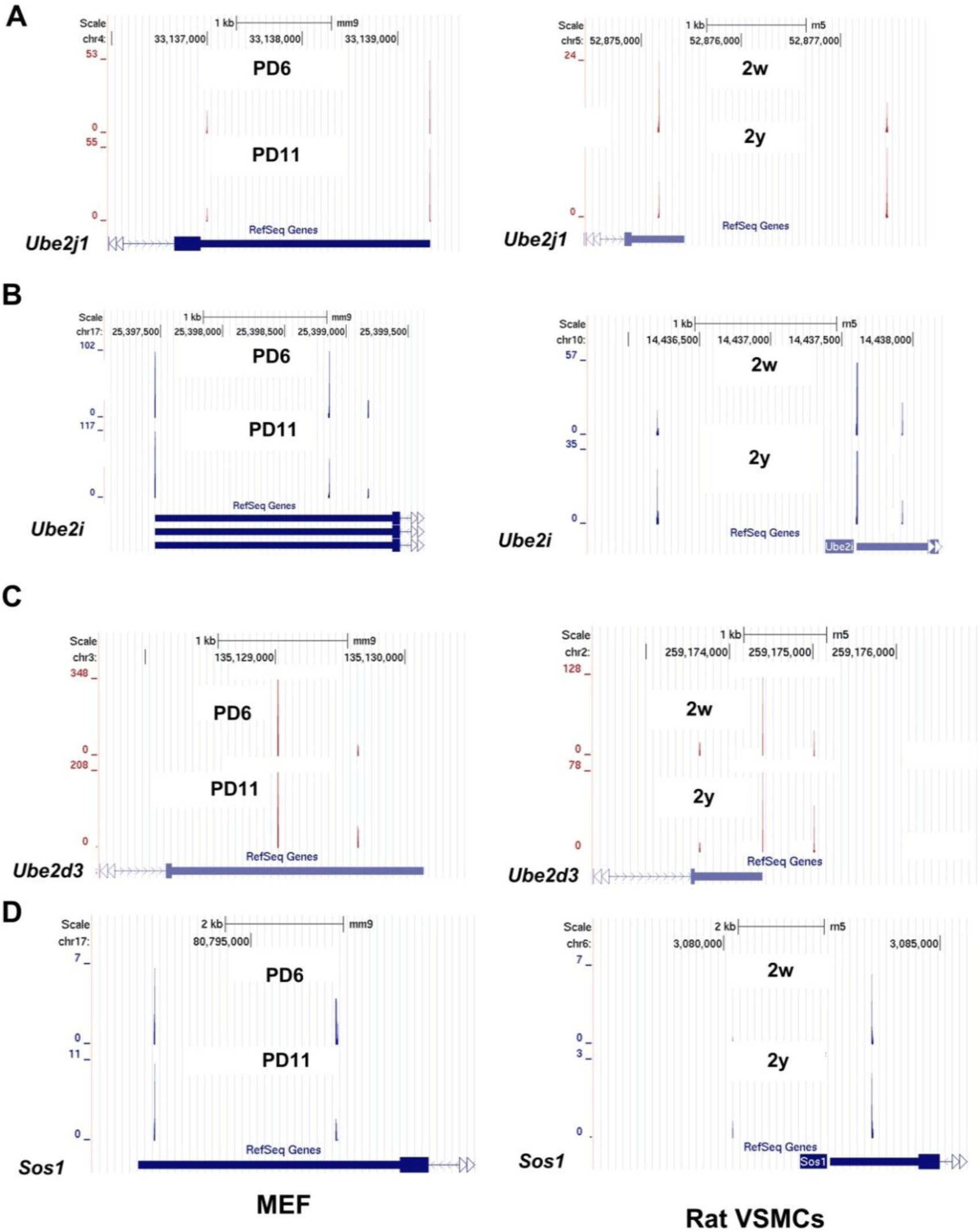
Examples of genes tended to use distal pAs in senescent MEFs and VSMCs of aged rat. (A-D) PA-seq tracks of *Ube2j1, Ube2i, Ube2d3,* and *Sos1* in senescent and young MEFs and VSMCs of old rat and young rat, respectively.

**Figure S9.**
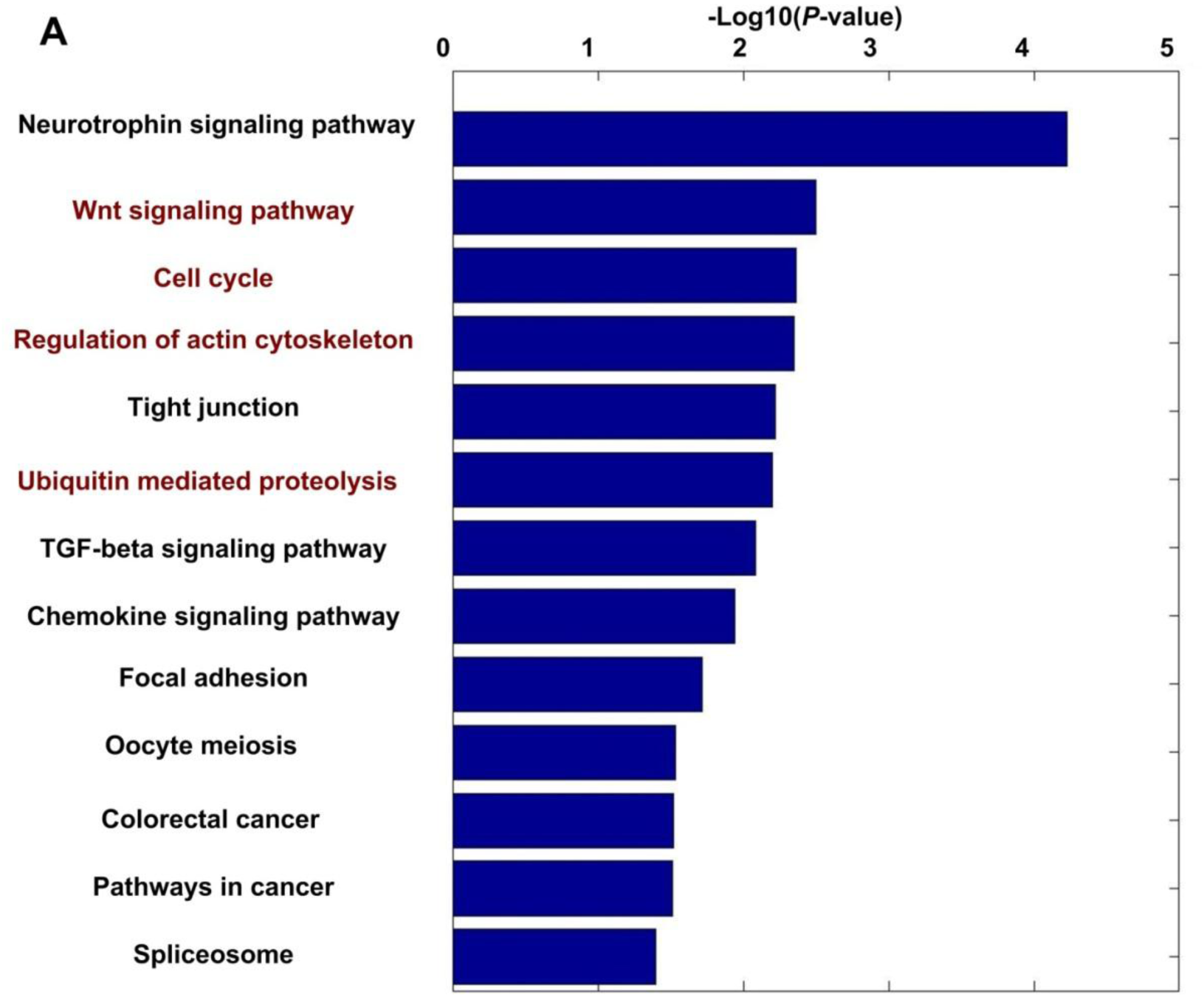
Genes progressively tended to use distal pAs during replicative senescence of MEFs are also enriched in senescence-related pathways. The common senescence-related pathways between Figure S9 and Figure5A are marked by red color.

**Figure S10.**
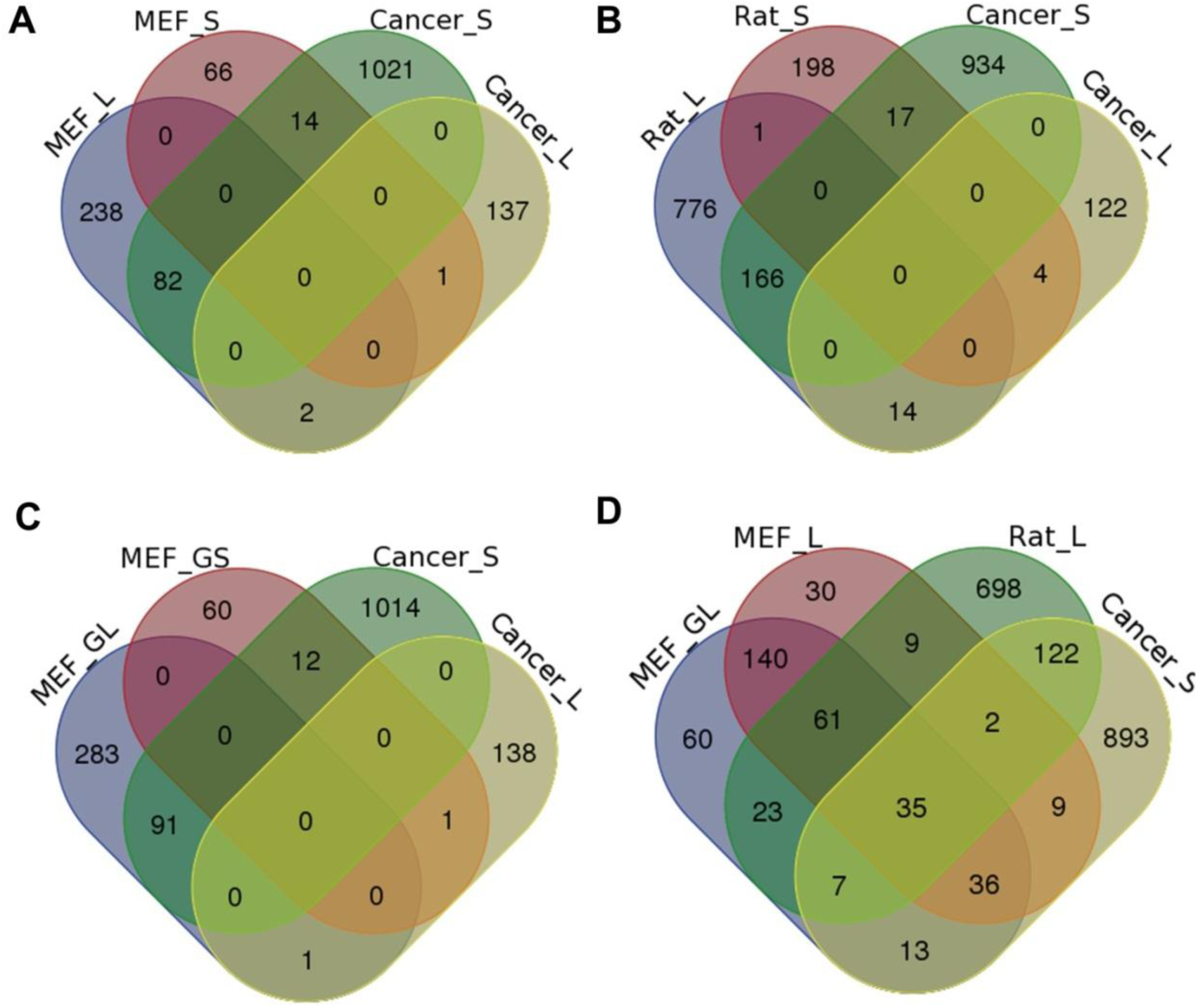
Comparison between genes significantly tended to use distal pAs and proximal pAs in senescent MEFs and VSMCs of old rat with genes significantly tended to use distal pAs and proximal pAs in cancer. (A) Venn diagram comparison among genes preferring distal (MEF_L) and proximal (MEF_S) pAs when comparing senescent MEFs (PD11) with Young MEFs (PD6) and genes preferring distal (Cancer_L) and proximal (Cancer_S) pAs when comparing tumors and normal tissues identified by Xia et al[3]. (B) Venn diagram comparison among genes preferring distal (Rat_L) and proximal (Rat_S) pAs when comparing VSMCs of old rat with young rat, Cancer_L and Cancer_S. (C) Venn diagram comparison among genes gradually preferred to use distal (MEF_GL) and proximal (MEF_GS) pAs during replicative senescence of MEFs, Cancer_L and Cancer_S. (D). Venn diagram comparison among MEF_L, Rat_L, MEF_GL and Cancer_S. MEF_L, MEF_S, Rat_L, Rat_S, MEF_GL and MEF_GS were identified by linear trend test with Benjamini-Hochberg (BH) false-discovery rate (FDR) at 5%.

**Figure S11.**
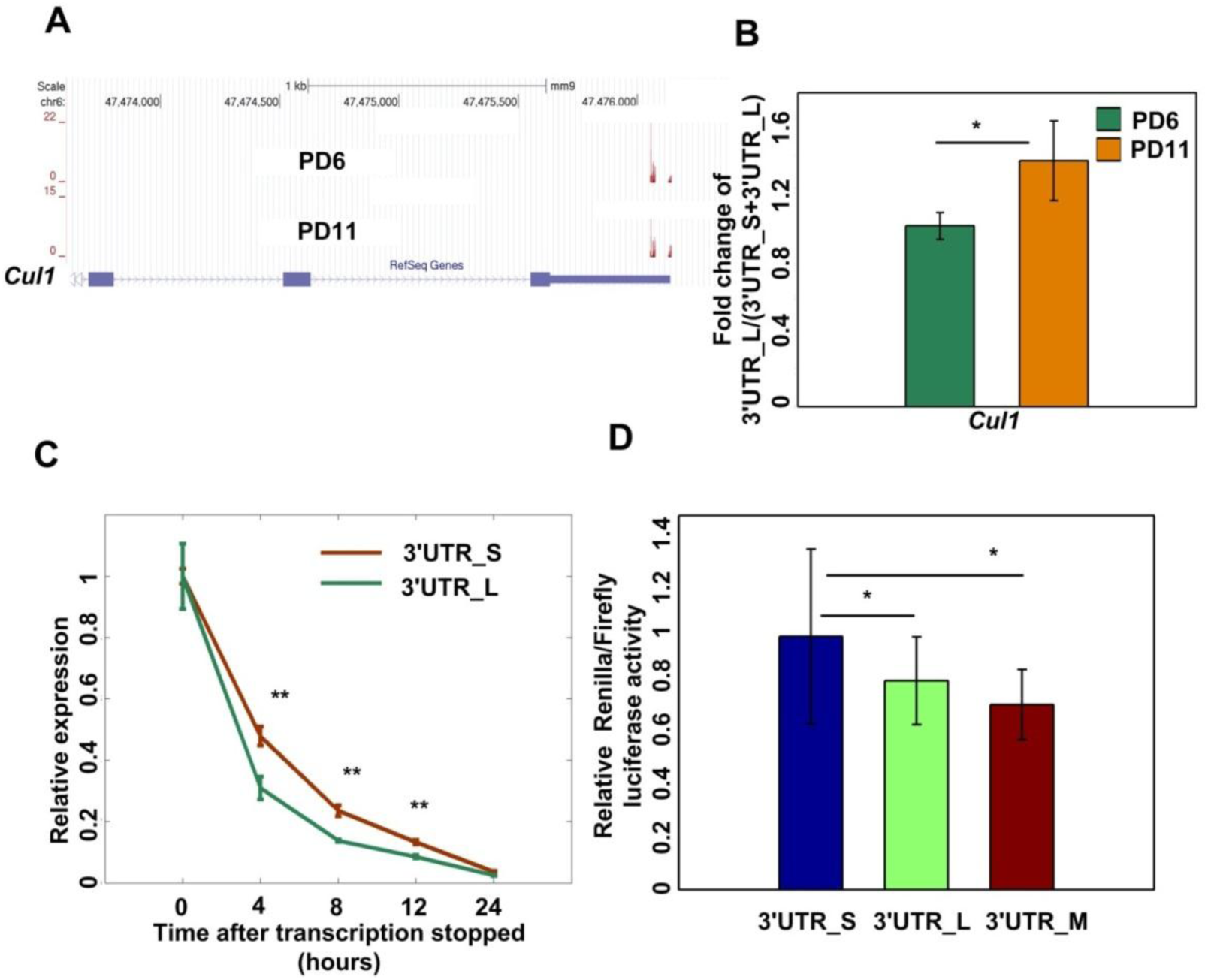
*Cull* with longer 3′ UTR has weaker stability and produces less protein. (A) PA-seq tracks of *Cull* in senescent and young MEFs. (B) qRT-PCR validation of pAs usage preference for *Cull* in senescent and young MEFs. (*) *P* <0.05, two tailed Wilcoxon signed rank test. (C) Comparing stability of the short 3′ UTR and the long 3′ UTR of *Cull* (named as 3′ UTR_L and 3′ UTR_S, respectively) in mouse fibroblast cell line NIH3T3. (**) *P* <0.01, two tailed Wilcoxon signed rank test. (D) Luciferase expression from a reporter containing the short 3′ UTR, compared to that from the reporter containing the long 3′ UTR and the long 3′ UTR with mutated PAS of proximal pA of *Cull* (named as 3′ UTR_L, 3′ UTR_S and 3′ UTR_M, respectively). (***) *P* <0.001, (**) *P* <0.01 and (*) *P* <0.05, two tailed Student’s t-test.

## SupplementaryTables

**Supplementary Table S1.** MEFs PA-seq reads mapping statistics.

| Sample | Total Reads | Strand specific reads | mappable Read1 | Read1 mapping rate | mappable Read2 | Read2 mapping rate |
| --- | --- | --- | --- | --- | --- | --- |
| G0 | 15,411,083 | 14,266,123 | 9,442,546 | 66.2% | 11,471,857 | 80.4% |
| PD6 | 9,423,643 | 7,784,321 | 4,478,586 | 57.5% | 5,689,182 | 73.1% |
| PD8 | 13,386,797 | 11,470,447 | 7,822,028 | 68.2% | 9,472,369 | 82.6% |
| PD10 | 8,779,338 | 7,307,736 | 3,818,641 | 52.3% | 4,799,825 | 65.7% |
| PD11 | 14,425,358 | 13,168,353 | 8,848,238 | 67.2% | 10,736,847 | 81.5% |

**Supplementary Table S2:** 18,639 Refined pAs identified in this study and their assigned categories (available as a separate Excel file).

**Supplementary Table S3:** MEFs RNA-seq reads mapping statistics.

| Sample | Total Reads | Strand specific reads | mappable Read1 | Read1 mapping rate | mappable Read2 |
| --- | --- | --- | --- | --- | --- |
| G0 | 15,809,150 | 13,997,188 | 88.5% | 15,809,150 | 88.6% |
| PD6 | 28,132,487 | 27,048,742 | 96.1% | 26,876,587 | 95.5% |
| PD8 | 25,659,880 | 24,321,086 | 94.8% | 24,175,499 | 94.2% |
| PD10 | 17,860,083 | 16,254,722 | 91.0% | 16,060,448 | 89.9% |
| PD11 | 16,396,313 | 14,101,035 | 86.0% | 13,980,835 | 85.3% |

**Supplementary Table S4:** Summary 3,165 genes with APA regulation (available as a separate Excel file).

**Supplementary Table S5.**
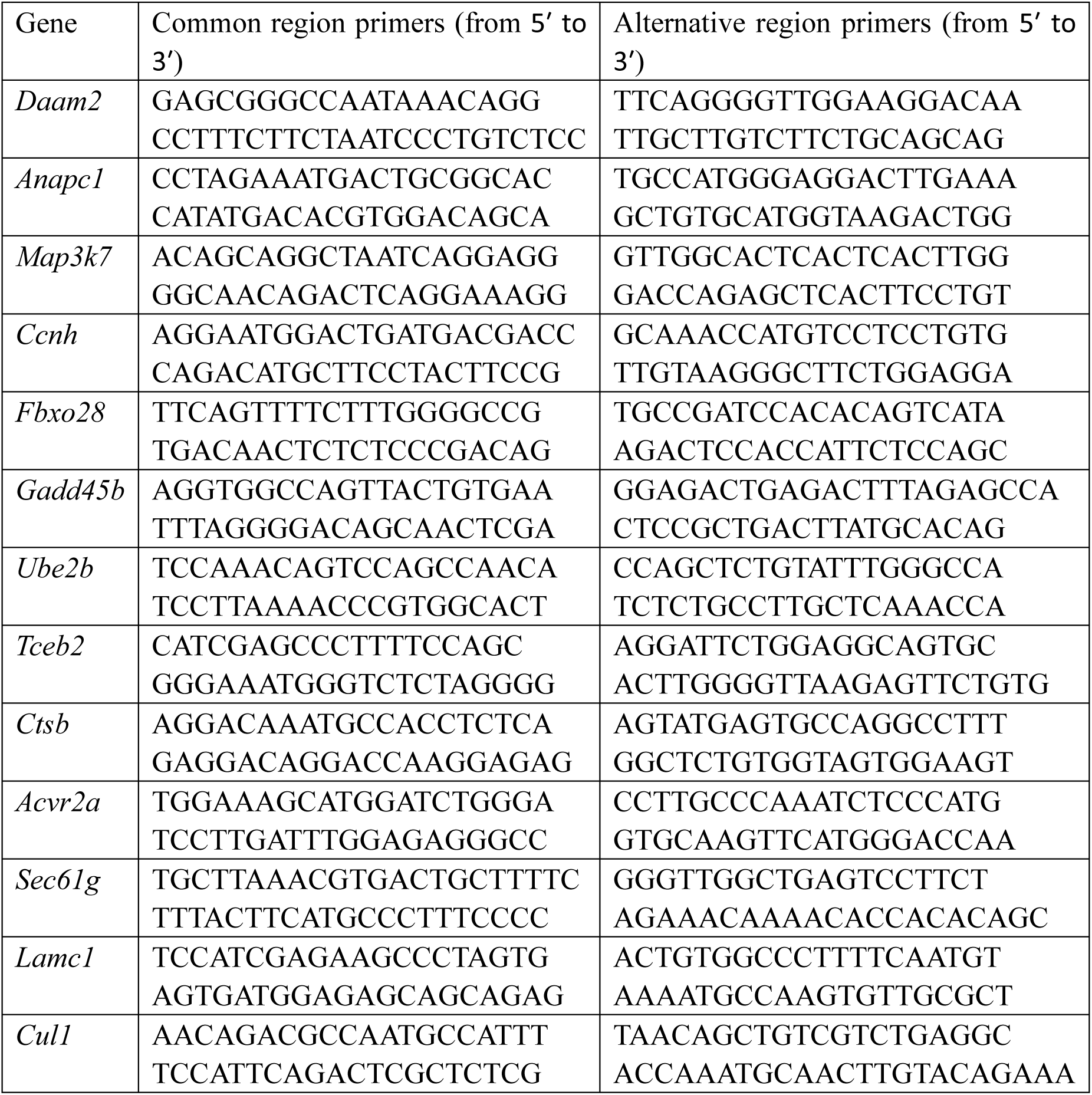
qRT-PCR primer sequences for genes be validated if they have significant APA usage switch.

**Supplementary Table S6.** Rat PA-seq reads mapping statistics.

| Sample | Total Reads | Strand specific reads | mappable Read1 | Read1 mapping rate | mappable Read2 | Read2 mapping rate |
| --- | --- | --- | --- | --- | --- | --- |
| 2 weeks | 26,122,246 | 20,468,514 | 8,818,888 | 43.1% | 11,694,741 | 57.1% |
| 2 years | 35,087,625 | 29,152,131 | 12,443,319 | 42.7% | 16,098,918 | 55.2% |

**Supplementary Table S7.** Gene list of genes with significant pA usage shift in replicative senescence of MEFs and aortic vascular smooth muscle cells of rats (VSMCs) with different ages. (Available as a separate Excel file).

**Supplementary Table S8.** Functional enrichment analysis for genes with significantly pA usage shift in replicative senescence of MEFs and aortic vascular smooth muscle cells of rats (VSMCs) with different ages. (Available as a separate Excel file).

**Supplementary Table 9.**
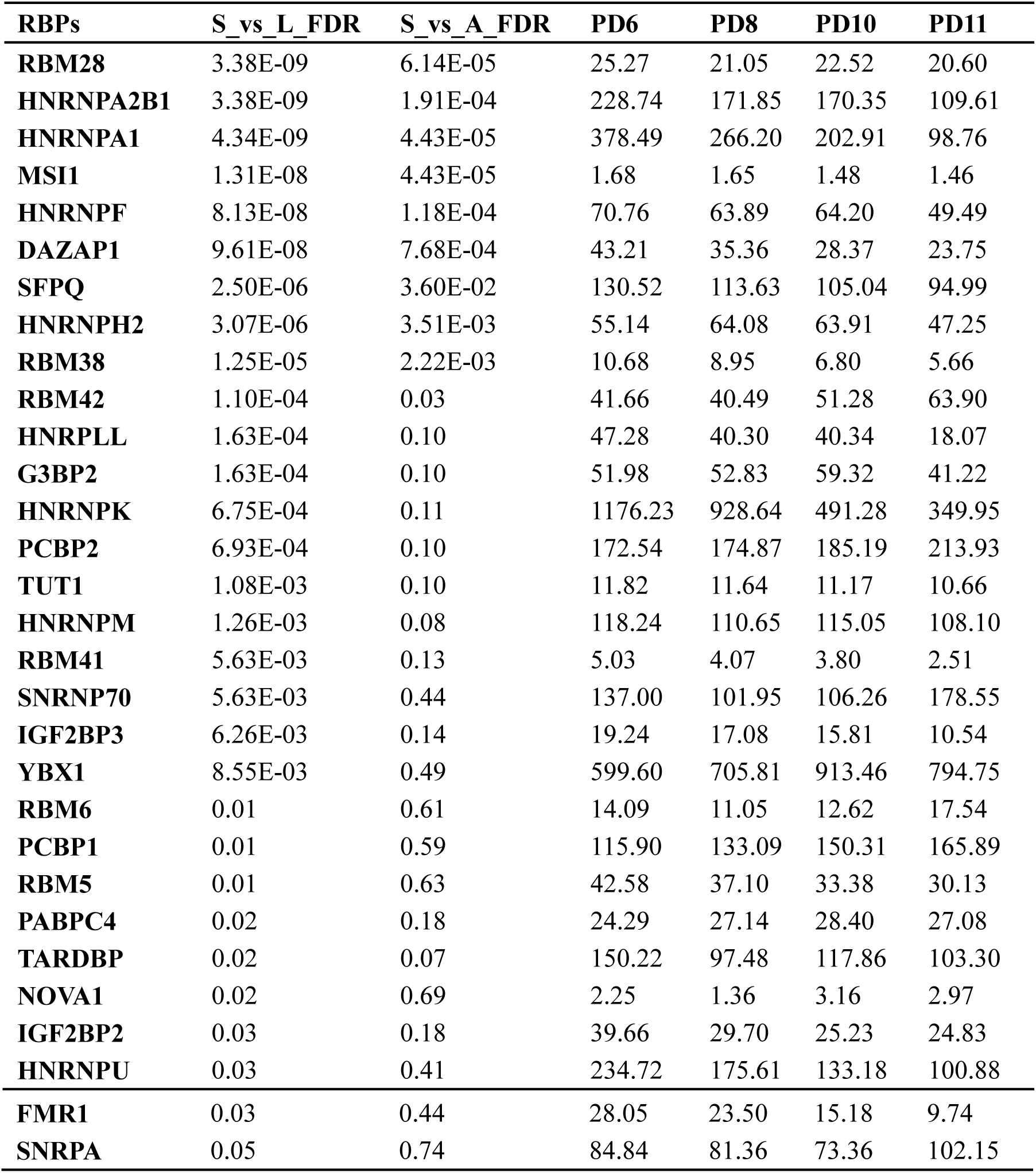
Summary of RBPs with higher binding site density in longer 3′ UTR than in shorter 3′ UTR regions. S_vs_L_FDR means the false discovery rate obtained by comparing the RNA binding site density between longer 3′ UTR and shorter 3′ UTR. S_vs_A_FDR means the false discovery rate obtained by comparing the RNA binding site density between longer 3′ UTR and alternative 3′ UTR. PD6, PD8, PD10, PD11 means the FPKM value of RBPs in the corresponding passage. Wilcox signed rank test was used to get the *P* value and Benjamini-Hochberg was used for multiple test correction.

**Supplementary Table10.**
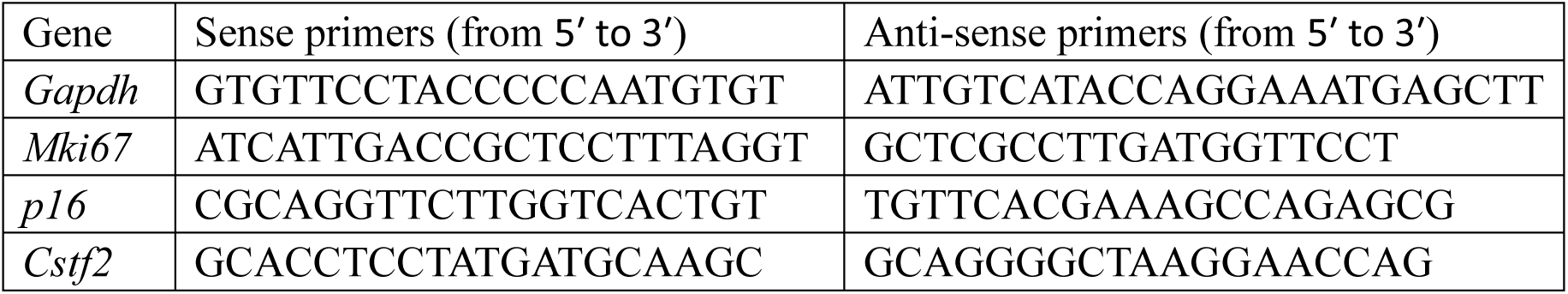
Primer sequences for genes which expression needed to be validated by qRT-PCR.

